# Musical prediction error responses similarly reduced by predictive uncertainty in musicians and non-musicians

**DOI:** 10.1101/754333

**Authors:** D.R. Quiroga-Martinez, N.C. Hansen, A. Højlund, M. Pearce, E. Brattico, P. Vuust

## Abstract

Auditory prediction error responses elicited by surprising sounds can be reliably recorded with musical stimuli that are more complex and realistic than those typically employed in EEG or MEG oddball paradigms. However, these responses are reduced as the predictive uncertainty of the stimuli increases. In this study, we investigate whether this effect is modulated by musical expertise. Magnetic mismatch negativity (MMNm) responses were recorded from 26 musicians and 24 non-musicians while they listened to low-and high-uncertainty melodic sequences in a musical multi-feature paradigm that included pitch, slide, intensity, and timbre deviants. When compared to non-musicians, musically trained participants had significantly larger pitch and slide MMNm responses. However, both groups showed comparable reductions of pitch and slide MMNm amplitudes in the high-uncertainty condition compared to the low-uncertainty condition. In a separate, behavioral deviance detection experiment, musicians were more accurate and confident about their responses than non-musicians, but deviance detection in both groups was similarly affected by the uncertainty of the melodies. In both experiments, the interaction between uncertainty and expertise was not significant, suggesting that the effect is comparable in both groups. Consequently, our results replicate the modulatory effect of predictive uncertainty on prediction error; show that it is present across different types of listeners; and suggest that expertise-related and stimulus-driven modulations of predictive precision are dissociable and independent.

## 1. Introduction

Prediction is fundamental for the perception of auditory sequences. When listening to a series of sounds, the brain generates expectations about future events partly based on the statistical regularities of the context and long-term knowledge of acoustic signals (Huron, 2006; Pearce, 2018). The violation of these expectations generates neural prediction error responses (den Ouden, Kok, & de Lange, 2012). So far, most research in this area has focused on very simple and artificial auditory contexts such as sequences of repeated tones or short tone patterns (Heilbron & Chait, 2018). As a consequence, little is known about how auditory prediction operates in more complex, real-world settings.

In a previous study, we addressed this issue by measuring prediction error responses to surprising sounds embedded in auditory stimuli that resembled real music (Quiroga-Martinez et al., 2019). As a marker of prediction error, we recorded the magnetic counterpart of the mismatch negativity (MMNm), which is a well-studied brain response to sounds that violate auditory regularities (Garrido, Kilner, Stephan, & Friston, 2009; Näätänen, Gaillard, & Mäntysalo, 1978). We compared a low-uncertainty condition —referred to as low-entropy or LE—which consisted of a simple and repetitive pitch pattern, with a high-uncertainty condition—referred to as high-entropy or HE—which consisted of more realistic and less predictable non-repetitive melodies. Note that entropy was used as a measure of uncertainty. Pitch, intensity, timbre and slide (i.e. pitch glide) violations were introduced. We found reliable MMNm responses to the violations in both conditions, thus demonstrating that low-level prediction error responses could be elicited in a constantly changing and more ecologically valid auditory stream.

Interestingly, even though MMNm responses were reliable, their amplitudes were reduced in the HE context compared to the LE context, for pitch and slide deviants. This is consistent with predictive processing theories which propose that prediction error responses are reduced in contexts with high as compared to low uncertainty or, equivalently, low as compared to high precision (Clark, 2016; Feldman & Friston, 2010; Hohwy, 2013; Ross & Hansen, 2016; Vuust, Dietz, Witek, & Kringelbach, 2018). The ensuing precision-weighted prediction error would ensure that primarily reliable sensory signals drive learning and behavior. While a growing body of research already provides evidence for this phenomenon in the auditory modality (Garrido, Sahani, & Dolan, 2013; Hsu, Bars, Hämäläinen, & Waszak, 2015; Lumaca, Haumann, Brattico, Grube, & Vuust, 2019; Sedley et al., 2016; Sohoglu & Chait, 2016; Southwell & Chait, 2018), our study was the first to show its presence in a more ecologically valid setting such as music listening. Furthermore, the findings also pointed to a feature-selective effect in which only prediction error responses related to the manipulated auditory feature—pitch, in our case—are modulated by uncertainty.

In the present work, we elaborate on this finding and investigate whether the effect of uncertainty on auditory prediction error is modulated by musical expertise. This question is motivated by research showing that musicians tend to exhibit stronger auditory prediction error responses than non-musicians. For example, larger MMN responses are often found for musically trained subjects, especially for pitch-related deviants (Brattico et al., 2009; Fujioka, Trainor, Ross, Kakigi, & Pantev, 2004; Koelsch, Schröger, & Tervaniemi, 1999; Putkinen, Tervaniemi, Saarikivi, Ojala, & Huotilainen, 2014; Tervaniemi, Huotilainen, & Brattico, 2014; Vuust, Brattico, Seppänen, Näätänen, & Tervaniemi, 2012; Vuust et al., 2005). This has led some to propose that musical training enhances the precision of auditory predictive models (Hansen & Pearce, 2014; Hansen, Vuust, & Pearce, 2016; Vuust et al., 2018), as more precise representations of musically relevant regularities would facilitate the detection of unexpected sounds.

Crucially, a distinction can be made between expertise-driven and stimulus-driven precision or uncertainty. The former corresponds to the fine-tuning of predictive models by musical training, whereas the latter refers to the uncertainty inferred from the stimulus currently being listened to. Note that stimulus-driven uncertainty was the one manipulated in Quiroga-Martinez et al., (2019). Consequently, our goal here is to address whether its effect on prediction error is modulated by expertise-driven precision. Thus, we conjectured that when musical sequences are predictable, long-term knowledge of music would only have a moderate impact on the processing of sounds. Conversely, when musical stimuli become more unpredictable, listeners would need to rely more on their musical knowledge, which would provide a greater processing advantage to musically trained participants. Therefore, we hypothesized an interaction effect in which the modulation of prediction error by uncertainty would be less pronounced for musicians than for non-musicians.

In this study, we used magnetoencephalography (MEG) and behavioral measures to test this hypothesis, employing the same stimuli and experimental designs as in Quiroga-Martinez et al. (2019). To this purpose, we compared a group of musicians with the group of non-musicians reported in the previous study. In the MEG experiment, participants passively listened to high-and low-entropy melodic sequences where pitch, intensity, timbre and slide deviants were introduced. In the behavioral experiment, participants were asked to detect pitch deviants embedded in different melodies and report the subjective confidence in their responses. In this case, five levels of context uncertainty were employed in order to detect fine-grained effects of predictive precision and dissociate pitch-alphabet size—the number of pitch categories used in the melodies—and repetitiveness as sources of uncertainty. We expected musicians to exhibit smaller reductions in MMNm responses, deviance detection scores, and confidence ratings than non-musicians, as the uncertainty of auditory contexts increased. Finally, we performed source reconstruction on the MMNm responses and the difference in MMNm amplitude between HE and LE conditions in order to further our understanding of the neural underpinnings of the precision-weighting effect.

## 2. Method

For a more detailed description of the methods, please see Quiroga-Martinez et al., (2019). The data, code and materials necessary to reproduce these experiments and results are openly available at: http://bit.ly/music_entropy_MMN

### 2.1. MEG experiment

#### 2.1.1. Participants

Twenty-six musicians and twenty-four non-musicians participated in the experiment (see Table 1 for demographics). The non-musicians’ group was the same as the one reported in Quiroga-Martinez et al., (2019). All participants were right-handed with no history of neurological conditions, and did not possess absolute pitch. The musical training subscale of the Goldsmiths Musical Sophistication Index (GMSI) was used as a self-report measure of musical expertise (Müllensiefen, Gingras, Musil, & Stewart, 2014) and both the melody and rhythm parts of the Musical Ear Test (MET) were used as objective measures of musical skills (Wallentin, Nielsen, Friis-Olivarius, Vuust, & Vuust, 2010). GMSI values (t = 17.9, p < .001) and MET total values (t = 5.2, p < .001) were significantly higher for musicians than for non-musicians. Participants were recruited through an online database and agreed to take part in the experiment voluntarily. All participants gave informed consent and were paid 300 Danish kroner (approximately 40 euro) as compensation. Data from two musicians (not included in the reported demographics) were excluded from the analysis due to artefacts related to dental implants. The study was approved by the Central Denmark Regional Ethics Committee (De Videnskabsetiske Komitéer for Region Midtjylland in Denmark) and conducted in accordance with the Helsinki declaration.

**Table 1.** Participants’ demographic and musical expertise information in the two experiments. M = musicians, NM = non-musicians, Beh = behavioral.

| Experiment | Group | Age (mean) | Age (SD) | Female | Male | GMSI (mean) | GMSI (SD) | MET Mel (mean) | MET Mel (SD) | MET Rhy (mean) | MET Rhy (SD) | MET Total (mean) | MET Total (SD) |
| --- | --- | --- | --- | --- | --- | --- | --- | --- | --- | --- | --- | --- | --- |
| MEG | NM (n=24) | 26.54 | 3.4 | 13 | 11 | 10.17 | 3.48 | 33.17 | 5.39 | 35.79 | 5.33 | 69.12 | 9.44 |
|  | M (n=26) | 24.15 | 2.89 | 10 | 16 | 35.85 | 6.35 | 41.5 | 4.43 | 40.77 | 4.55 | 82.27 | 8.35 |
| Beh | NM (n=21) | 21.9 | 5.18 | 16 | 5 | 12.9 | 5.77 | - | - | - | - | - | - |
|  | M (n=24) | 22.75 | 4.42 | 14 | 10 | 35.42 | 5.69 | - | - | - | - | - | - |

#### 2.1.2. Stimuli

Low-entropy (LE) and high-entropy (HE) conditions were included in the experiment. LE stimuli corresponded to a simple four-note repeated pitch pattern known as the Alberti bass, which has previously been used in musical MMNm paradigms (Vuust et al., 2011; Vuust, Liikala, Näätänen, Brattico, & Brattico, 2016). In contrast, HE stimuli consisted of a set of major and minor versions of six novel melodies which did not have a repetitive internal structure and spanned a broader local pitch range than LE stimuli (Figure 1; see Supplementary file 1 in Quiroga-Martinez et al., 2019 for the full stimulus set). Individual HE and LE melodies were 32-notes long, lasted eight seconds, and were pseudorandomly transposed from 0 to 5 semitones upwards. The order of appearance of the melodies was pseudorandom. After transposition during stimulation, the pitch-range of the HE condition spanned 31 semitones from B3 (F_0_ ≈ 247 Hz) to F6 (F_0_ ≈ 1397 Hz). LE melodies were transposed to two different octaves to cover approximately the same pitch range as HE melodies. The uncertainty of the stimuli was estimated with Information Dynamics of Music (IDyOM), a variable-order Markov model of auditory expectation (Pearce, 2005, 2018). When predicting pitch continuations based on a training corpus of Western tonal hymns and folk songs, this model confirmed higher mean entropy (which is a measure of uncertainty) and information content (which is a measure of surprise) for HE as compared to LE melodies (see Quiroga-Martinez et al., 2019 for more details).

**Figure 1.**
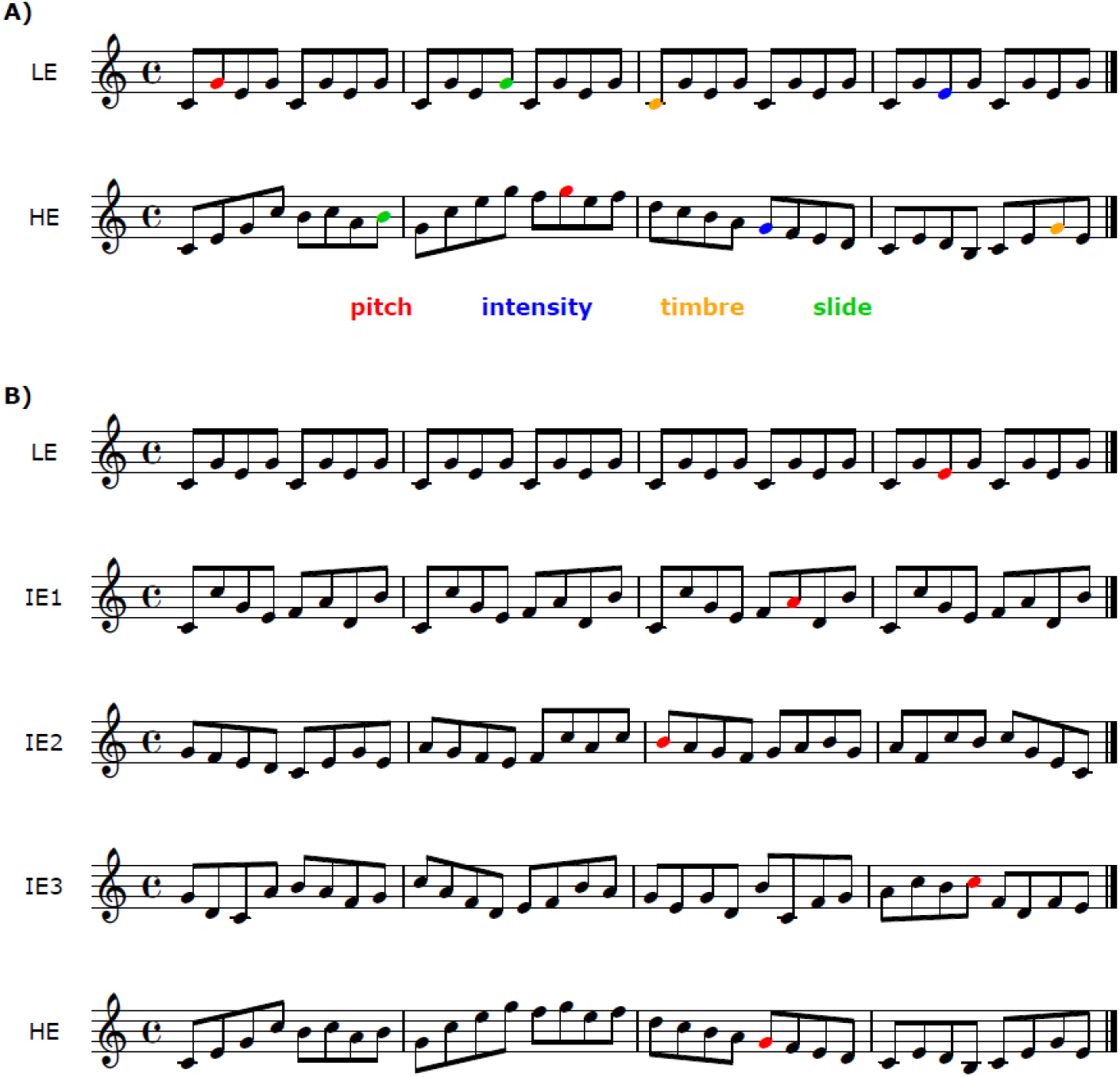
Examples of the individual melodies employed in A) the MEG and B) the behavioral experiment. Colored notes represent deviants. LE = low entropy, IE = intermediate entropy, HE = high entropy. For the full stimulus set see Quiroga-Martinez et al. (2019) and the online repository.

For stimulus delivery, a pool of 31 standard piano tones was created with the “Warm-grand” sample in Cubase (Steinberg Media Technology, version 8). Each tone was 250 ms long, was peak-amplitude normalized and had 3-ms-long fade-in and fade-out to prevent clicking. No gaps between tones were introduced. For the creation of deviants, the standards were modified as follows. Pitch: +50 cents; intensity: −20 dB; timbre: band-pass filter (1-4 kHz); slide: continuous pitch glide from −2 semitones. Deviants were created with Audition (Adobe Systems Incorporated, version 8).

Each condition was presented in a separate group of three consecutive blocks. Within each block, melodies were played one after the other without pauses. At the beginning of each block, a melody with no deviants was added to ensure a certain level of auditory regularity at the outset. One deviant per feature was introduced in each melody. There were 144 deviants per feature in each condition. The position of each deviant was defined by segmenting the melody in groups of four notes, selecting some of these groups, and choosing randomly any of the four places within a group, with equal probability. The order of appearance of the different types of deviants was pseudorandom, so that no deviant followed another deviant of the same feature. The selection of four-note groups was counterbalanced among trials attending to the constraints of a combined condition included to assess the predictive processing of simultaneous musical streams (see Quiroga-Martinez et al., 2019 for further details). The analysis of the combined condition is beyond the scope of this article and will be presented elsewhere. HE and LE conditions were counterbalanced across participants and always came after the combined condition.

#### 2.1.3. Procedures

Participants gave their consent after receiving oral and written information, and then completed the MET, filled out the GMSI questionnaire and put on MEG-compatible clothes. Electrodes and HPI coils were attached to their skin and their heads were digitized. During the recording, participants were sitting upright in the MEG scanner looking at a screen. Before presenting the musical stimuli, their individual hearing threshold was measured through a staircase procedure and the sound level was set at 60dB above threshold. Participants were instructed to watch a silent movie of their choice, ignore the sounds and move as little as possible. They were told there would be musical sequences playing in the background interrupted by short pauses so that they could take a break and adjust their posture. Sounds were presented through isolated MEG-compatible ear tubes (Etymotic ER•30). The recording lasted approximately 90 minutes and the whole experimental session took between 2.5 and 3 hours including consent, musical expertise tests, preparation, instructions, breaks, and debriefing.

#### 2.1.4. MEG recording and analyses

Brain magnetic fields were recorded with an Elekta Neuromag MEG TRIUX system with 306 channels (204 planar gradiometers and 102 magnetometers) and a sampling rate of 1000 Hz. Continuous head position information (cHPI) was obtained with four coils (cHPI) attached to the forehead and the mastoids. Offline, the temporal extension of the signal source separation (tSSS) technique (Taulu & Simola, 2006) was used to isolate signals coming from inside the skull employing Elekta’s MaxFilter software (Version 2.2.15). This procedure included movement compensation for all participants except two non-musicians, for whom continuous head position information was not reliable due to suboptimal placement of the coils. These participants, however, exhibited reliable auditory event-related fields (ERFs), as successfully verified by visually inspecting the amplitude and polarity of the P50(m) component. Electrocardiography, electrooculography, and independent component analysis were used to correct for eye-blink and heartbeat artifacts, employing a semi-automatic routine (FastICA algorithm and functions “find_bads_eog” and “find_bads_ecg” in MNE-Python) (Gramfort, 2013). Visual inspection of the rejected components served as a quality check.

Using the Fieldtrip toolbox (version r9093) (Oostenveld, Fries, Maris, & Schoffelen, 2011) in MATLAB (R2016a, The MathWorks Inc., Natick, MA), epochs comprising a time window of 400 ms after sound onset were extracted and baseline-corrected with a pre-stimulus baseline of 100 ms. Epochs were then low-pass filtered with a cut-off frequency of 35 Hz and down-sampled to a resolution of 256 Hz. For each participant, ERFs were computed by averaging the responses for all deviants for each feature and averaging a selection of an equal number of standards. These were selected by finding, for each single deviant, a standard tone that was not preceded by a deviant and was in the same position of the same HE or LE melody—although not necessarily the same transposition—in a different trial. This ruled out artefacts related to the difference in noise between conditions—since there are many more standards than deviants— and the position of the deviant within the melody. After averaging, planar gradiometers were combined by computing root mean square values. Finally, a new baseline correction was applied and MMNm difference waves were computed by subtracting the ERFs of standards from the ERFs of deviants.

Statistical analyses were performed on combined gradiometer data. For the main analyses, a mass univariate approach was used in combination with cluster-based permutations (Maris & Oostenveld, 2007) for family-wise error correction. Two-sided paired-and independent-samples t-tests were used for within-and between-subjects contrasts, respectively. The cluster-forming alpha level was .05, the cluster-level statistic was the maximum sum of *t*-values *(maxsum)* and the number of permutations was set to 10,000. All tests were conducted for each feature separately in a time window between 100 and 250 ms, which covers the typical latency of the MMN (Näätänen, Paavilainen, Rinne, & Alho, 2007). To assess the elicitation of the

MMNm, we compared the ERFs of standards with the ERFs of deviants for each group independently. The main effect of entropy was assessed by comparing, for each feature, the MMNm responses of all participants for LE and HE conditions. The main effect of expertise was assessed by comparing the average of LE and HE responses between groups. The entropy-by-expertise interaction was tested by subtracting HE from LE MMNm responses for each participant, and comparing the resulting differences between groups. Post-hoc, exploratory tests of simple effects were performed for the effect of entropy and expertise for each group and condition, respectively.

To assess the relative evidence for the null and alternative hypotheses, a secondary Bayesian analysis was performed on mean gradient amplitudes (MGA), which were obtained as the mean activity ±25 ms around the MMNm peak, defined as the highest local maxima of the ERF between 100 and 250 ms after sound onset. This average was obtained from the four temporal combined gradiometers in each hemisphere with the largest P50(m) response (right channels: 1342-1343, 1312-1313, 1322-1323, 1332-1333; left channels: 0222-0223, 0212-0213, 0232-0233, 0242-0243). Using R (R Core Team, 2019), the differences between HE and LE MMNm amplitudes were computed for each participant and used as the dependent variable in a Bayesian mixed-effects model including parameters for the effects of feature, hemisphere and group and their interactions (brms package, Bürkner, 2017). Participants were included as a random effect with respect to the intercept and the slopes of feature and hemisphere. Priors were taken from our previous work with the non-musicians’ group (see the analysis scripts and saved model fits in the online repository for a full description of priors and parameters). For the effect of expertise and the interactions with hemisphere and feature, a conservative prior was set with a mean of 0 and a standard deviation of 3 fT/cm, which is around half of the effect of entropy for the pitch MMNm in non-musicians. This prior assumes that small effect modulations are most likely and that situations in which the effect of entropy in musicians disappears, changes direction, or is at least twice the effect in non-musicians are unlikely. Inference was based on 95% credible intervals, Bayes Factors (BF) and posterior probabilities, as estimated for each feature and hemisphere (“hypothesis” function, *brms* package).

#### 2.1.5. Source reconstruction

Source reconstruction was performed with the Multiple Sparse Priors (MSP) method (K. J. Friston et al., 2008) implemented in SPM12 (version 7478). Only data from twenty musicians and twenty non-musicians were included, since individual anatomical magnetic resonance images (MRI) were available for these participants only. For one of the excluded musicians and one of the excluded non-musicians the images were corrupted by artefacts, whereas the remaining excluded participants did not attend the MRI session. Brain scans were obtained with a Magnetization-prepared two rapid gradient echo (MP2RAGE) sequence (Marques et al., 2010) in a Siemens Magnetom Skyra 3T scanner, which produced two images that were combined and motion-corrected to form unified brain volumes. These volumes were segmented, projected into MNI coordinates, and automatically coregistered with the MEG sensor positions using digitized head shapes and preauricular and nasion landmarks. Coregistration outputs were visually inspected. Lead-fields were constructed using a single-shell BEM model with 20.484 dipoles (fine grid). A volume of the inverse solution was created for each participant feature and condition, in the following time windows: 175-215 ms for pitch, 110-150 ms for timbre and intensity, and 275-315 ms for slide. These time windows were chosen based on the peak MMNm amplitudes for each feature. Source reconstruction was also conducted for the differences between HE and LE conditions for pitch and slide MMNm amplitudes, with the aim to reveal the neural substrates of the entropy effect. To this end, individual volumes of the inverse solution were obtained for a time window between 150 and 200 ms. Note that the case of the slide deviant is somewhat particular since the peak MMNm response occurred later than expected, whereas the effect of entropy was restricted to an earlier time window. For greater detail and an interpretation of this result, see Quiroga-Martinez et al. (2019). The volumes for each feature and condition, as well as the volumes for the entropy effect, were submitted to a one-sample t-test to reveal the sources consistently identified across all participants. The error rate of voxel-wise multiple tests was corrected with random field theory with a cluster-level alpha threshold of 0.05 (Worsley, 2007).

### 2.2. Behavioral experiment

#### 2.2.1. Participants

Twenty-four musicians and twenty-one non-musicians participated in the behavioral experiment (Table 1). The non-musicians’ group is the same as the one reported in Quiroga-Martinez et al. (2019). Musical expertise was measured with the GMSI musical training subscale which yielded significantly higher scores for musicians than for non-musicians (t = 13.14, p < .001). Participants were recruited through an online database for experiment participation, agreed to take part voluntarily, gave their informed consent and received 100 Danish kroner (approximately 13.5 euro) as compensation. The data from all participants were analyzed, since above-chance deviance detection was verified in all cases. The sample size was chosen to be comparable to that of the MEG experiment. Two musicians and two non-musicians had previously participated in the MEG experiment.

#### 2.2.2. Experimental design

Five conditions were included (Figure 1). Two of them correspond to the HE and LE conditions of the MEG experiment and employ a selection of the respective melodies. Three additional conditions with intermediate levels of entropy (IE1, IE2, IE3) were included to investigate whether more fine-grained manipulations of uncertainty modulate prediction error responses in musicians, as was previously shown in non-musicians. The pitch alphabet of these conditions spanned eight tones and was always the same, comprising a major diatonic scale from C4 to C5. Note that, in the MEG experiment, HE stimuli were not only less repetitive but also had a larger pitch alphabet than LE (at least before transposition during the experiment). In contrast, in the IE conditions we manipulated uncertainty by changing repetitiveness only. Thus, IE1 consisted of a repeated eight-note pattern, IE2 consisted of proper melodies with less constrained repetition, and IE3 consisted of pseudorandom orderings of the tones. Note that the contrast LE > IE1 would reveal whether the pitch-alphabet size alone is sufficient to modulate prediction error, whereas the comparisons IE1 > IE3, IE1 > IE2 and IE2 > IE3 would reveal the same with regard to repetitiveness.

For each single melody in the experiment, a target version was created by raising the pitch of a tone by 25 cents. This deviation was smaller than in the MEG experiment to avoid ceiling effects observed for non-musicians during piloting. The target tone was located in a random position in the second half of each melody. All melodies were 32-notes long and were played with the same sound pool as the MEG experiment. There were ten target melodies and ten foil melodies (with no deviants) per condition. Participants were instructed to listen to the melodies, decide after each of them whether an out-of-tune note was present or not, and report how certain they were about their answer on a scale from 1 (not certain at all) to 7 (completely certain). The experimental session lasted around 30 minutes.

#### 2.2.3. Statistical analyses

We used signal detection theory to analyze accuracy (Macmillan, 2004), based on the assumption that larger prediction error responses would enhance the ability to distinguish target from non-target stimuli. For each condition, d-prime (*d*’) scores were computed as a measure of sensitivity and criterion (c) scores were computed as a measure of response bias. In the few cases where participants achieved 100% or 0% of hits or false alarms, values were adjusted to 95% and 5% respectively, to avoid infinite values in the estimations (Macmillan, 2004).

Statistical analyses were run in R. For *d’* scores, different mixed-effects models were estimated using maximum likelihood (“lmer” function, lme4 package, Bates, Mächler, Bolker, & Walker, 2015), and compared using likelihood ratio tests and Akaike Information Criteria (AIC). Model *d0* included only an intercept as a fixed effect, whereas two alternative models added categorical (*d1*) or continuous (*d2*) terms for the entropy conditions. For model *d2*, we assigned values 1, 2, 3, 4 and 5 to the conditions according to their estimated uncertainty, and treated them as a continuous linear predictor. This allowed us to assess the extent to which a linear decreasing trend was present in the data, as was done previously with non-musicians (Quiroga-Martinez et al., 2019). Building on these models, in *d1e* and *d2e* a term for musical expertise was added, and in *d1i* and *d2i* a term for the entropy-by-expertise interaction was further included. Random intercepts for participants were included in all models. Random slopes were not added, as the number of data points per participant was not sufficient to avoid overfitting. For *c* scores, mixed-effects models were similarly compared, including an intercept-only model (*cr0*), a model with a categorical effect of entropy (*cr1*), a model with an additional effect of expertise (*cr1e*) and a model with an additional term for the entropy-expertise interaction (*cr1i*). Random intercepts for participants were added.

Regarding confidence ratings, ordinal logistic regression was employed in the form of a cumulative-link mixed model (“clmm” function, ordinal package; Christensen, 2019) using logit (log-odds) as link. Models with an intercept only (co0), categorical (*co1s*) terms for entropy, and additional terms for expertise (*co1se*) and the entropy-by-expertise interaction (*co1si*) were estimated and compared. These models included random intercepts and slopes for participants. Unlike with *d’* scores, no model included continuous terms for entropy, since categorical models were previously shown to explain the data significantly better for nonmusicians. Moreover, note that the cumulative-link model estimates an intercept for each cut-point between adjacent categories in the response variable. Post-hoc, Bonferroni-corrected pairwise contrasts for the effect of entropy on confidence ratings, *d’* scores and *c* scores were conducted with the function “emmeans” (emmeans package; Lenth, Singmann, Love, Buerkner, & Herve, 2019) for musicians and non-musicians separately.

Bayesian estimation was used to assess the evidence for the entropy-by-expertise interaction. Models *d1i*, *d2i*, and *co1i* were re-estimated and labeled as *d1ib, d2ib* and *co1ib.* Priors were defined based on our previous work with non-musicians (see corresponding analyses scripts in the online repository for a full description of priors and parameters). For the continuous model of *d’* scores (*d2ib*), the prior for the entropy-by-expertise interaction was Gaussian with mean 0 and standard deviation 0.1, which corresponds to half of the slope for the effect of entropy previously estimated for non-musicians. This prior is conservative and implies that small effect modulations are deemed most likely, and that a complete absence, change of direction or excessive enhancement of the effect is considered unlikely. For the categorical model of *d’* scores (*d1ib*), a Gaussian prior with mean 0 and standard deviation 0.4 was used for each of the entropy conditions. This prior corresponds to about half of the difference between the LE and HE conditions and, as with the continuous model, is conservative and deems extreme modifications of the effect unlikely. Regarding confidence ratings, a similar conservative Gaussian prior was set for the interaction term, with mean 0 and standard deviation 0.35. This prior deems small effects as the most likely and odds modifications larger than twice (*e^2×0.35^ = 2*) or smaller than half (*e^2×-0·35^ = 0.5*) of the original effect as unlikely. Inference was based on 95% credible intervals, Bayes Factors and posterior probabilities, estimated for each feature and hemisphere (“hypothesis” function, *brms* package).

## 3. Results

### 3.1. Presence of the MMNm

As previously reported for the non-musicians (Quiroga-Martinez et al., 2019), we also found a difference between standards and deviants for each feature in the musicians’ group (all p < .001, supplementary file 1). This difference had virtually the same topography and polarity as previously reported, thus confirming the presence of the MMNm (Figure 2).

**Figure 2.**
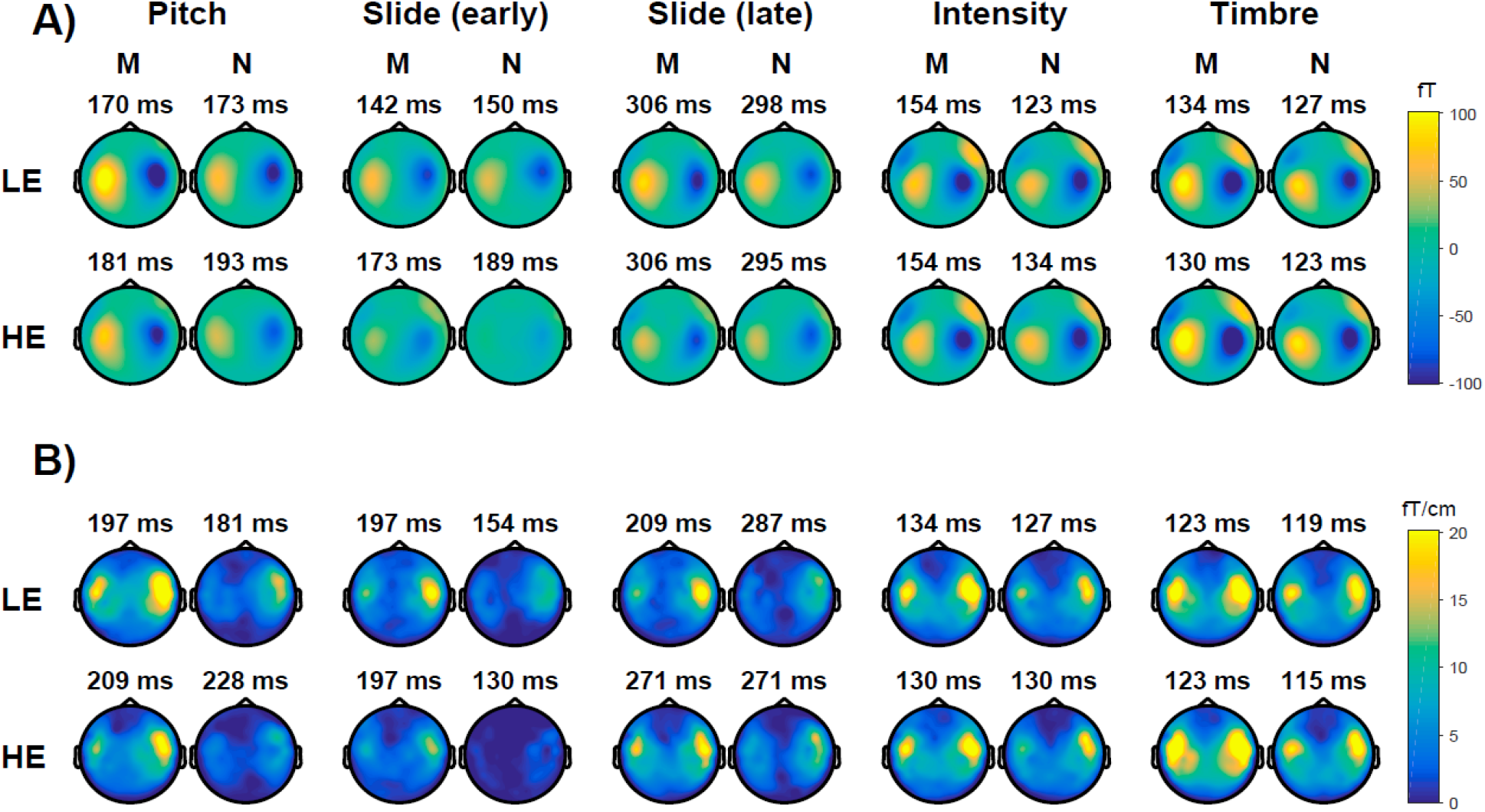
Topographic maps of the MMNm for all features, groups, and conditions in A) magnetometers and B) gradiometers. The activity corresponds to an average of ±25 ms around the peak latency, which is shown above each plot. The slide MMNm is displayed in both early and late time windows (see Quiroga-Martinez et al., 2019 for an explanation of early and late effects in the slide MMNm). LE = low entropy, HE = high entropy, M = musicians, N = non-musicians.

### 3.2. Effects of entropy, expertise, and interaction

There was a significant main effect of entropy for pitch (p < .001), slide (p < .001) and intensity (p < .001), but not for timbre (p = .068), in the MMNm responses. Analyses of simple effects revealed significant differences for pitch and slide in both groups, and for intensity in musicians only (Figure 3). A significant main effect of expertise was observed for pitch and slide, but not intensity or timbre, in the MMNm responses (Figure 4). The same pattern emerged when LE (pitch: p = .01; slide: p = .018; intensity: p = .99; timbre: p = .68) and HE (pitch: p = .005; slide: p = .014; intensity: p = .3; timbre: p = .89) conditions were analyzed separately. The entropy-by-expertise interaction was not significant for any of the four features (Figure 4).

**Figure 3.**
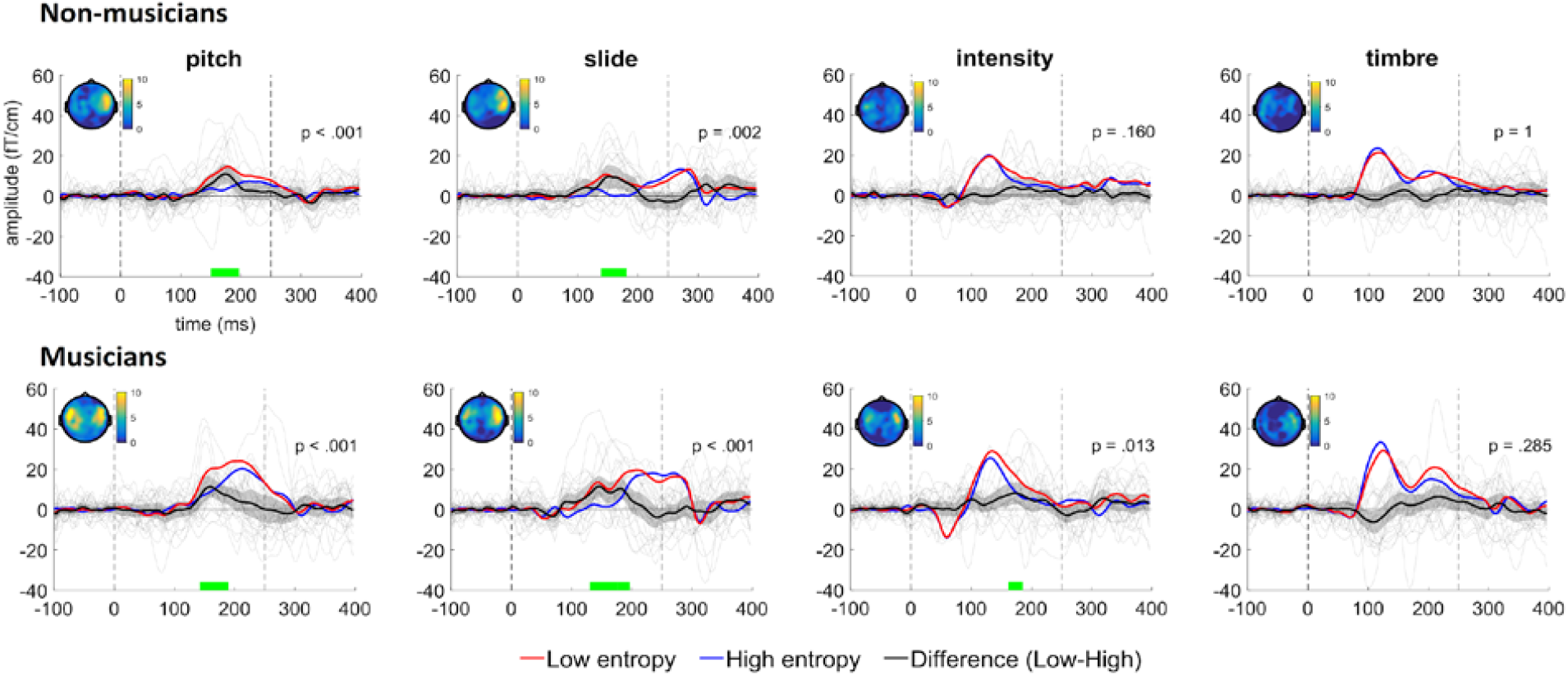
MMNm amplitudes for low-entropy (LE) and high-entropy (HE) conditions and the difference between conditions in both groups. The displayed activity corresponds to the average of the four right temporal combined gradiometers with the largest amplitude (channels 1342-1343, 1312-1313, 1322-1323 and 1332-1333). Gray lines depict individual MMNm responses. Shaded gray areas indicate 95% confidence intervals. Dashed vertical lines mark tone onsets. Topographic maps show activity ±25 ms around the peak difference. For descriptive purposes, green horizontal lines indicate when this difference was significant, according to the permutation tests. Note, however, that this is not an accurate estimate of the true extent of the effect (Sassenhagen & Draschkow, 2019).

**Figure 4.**
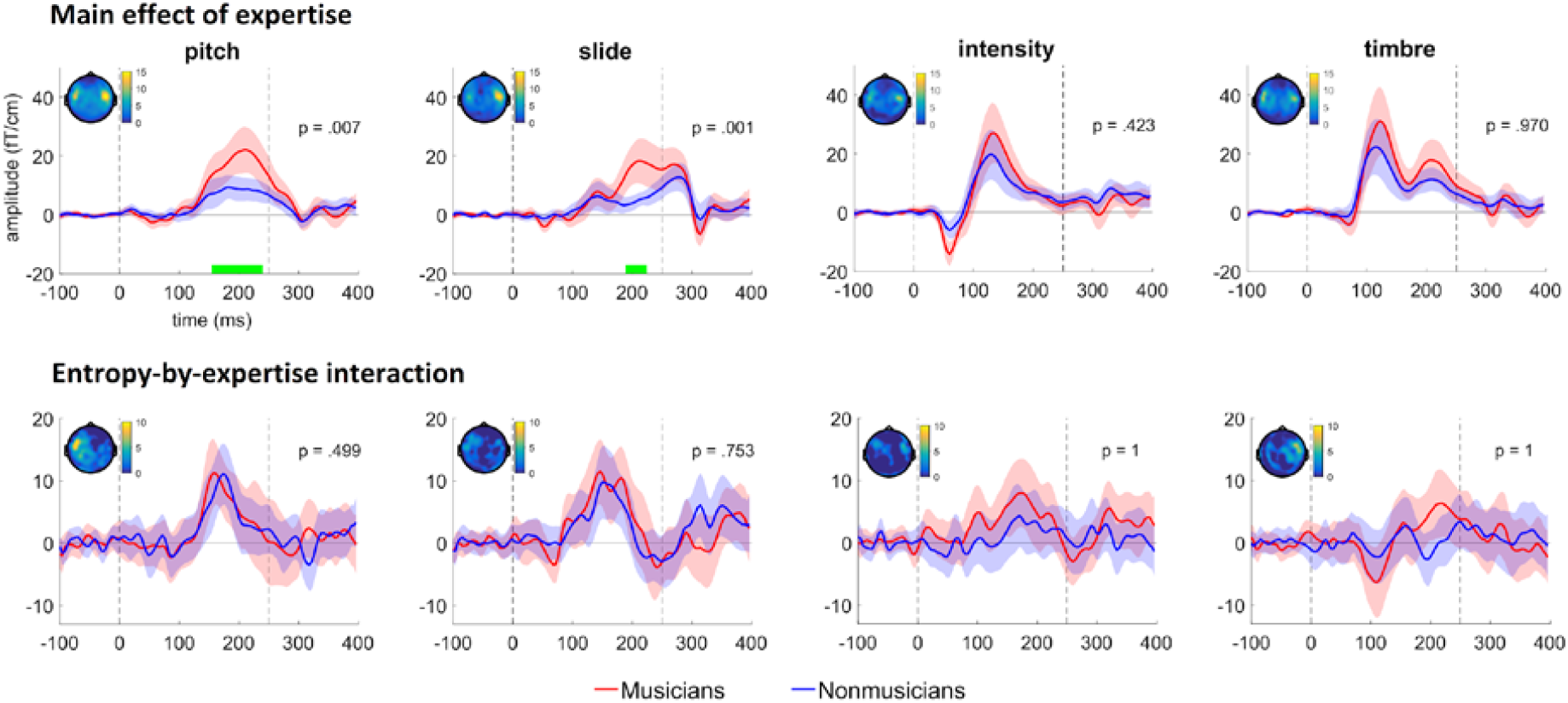
Activity related to the main effect of expertise and the entropy-by-expertise interaction—i.e. difference between low-entropy and high-entropy MMNm amplitudes for musicians and non-musicians. The displayed activity corresponds to the average of the four right temporal combined gradiometers with the largest amplitude (channels 1342-1343, 1312-1313, 1322-1323 and 1332-1333). Shaded areas indicate 95% confidence intervals. Dashed vertical lines mark tone onsets. Topographic maps show activity ±25 ms around the peak difference. For descriptive purposes, green horizontal lines indicate when this difference was significant. Note, however, that this is not an accurate estimate of the true extent of the effect (Sassenhagen & Draschkow, 2019).

Regarding secondary Bayesian analyses, the posterior distributions of the differences between musicians and non-musicians for each hemisphere and feature are shown in Figure 5b. 95% credible intervals included zero in all cases. Bayes factors suggested that the null hypothesis was between 1.18 to 3.06 times more likely than the alternative, and the posterior probability of a null effect varied between 0.62 and 0.75, depending on the feature and hemisphere. We regard this as anecdotal/inconclusive evidence for the null hypothesis. Moreover, Bayesian pairwise contrasts between features reproduced the patterns observed in the maximum likelihood estimates previously reported for non-musicians (Quiroga-Martinez et al., 2019), in which pitch and slide tended to have larger entropy-related reductions in MMNm amplitude than intensity and timbre, in the right but not the left hemisphere. For musicians this pattern was different, with conclusive evidence for a difference between pitch and timbre in both hemispheres, and moderate evidence for a difference between pitch and intensity in the left hemisphere (Table 2).

**Table 2.**
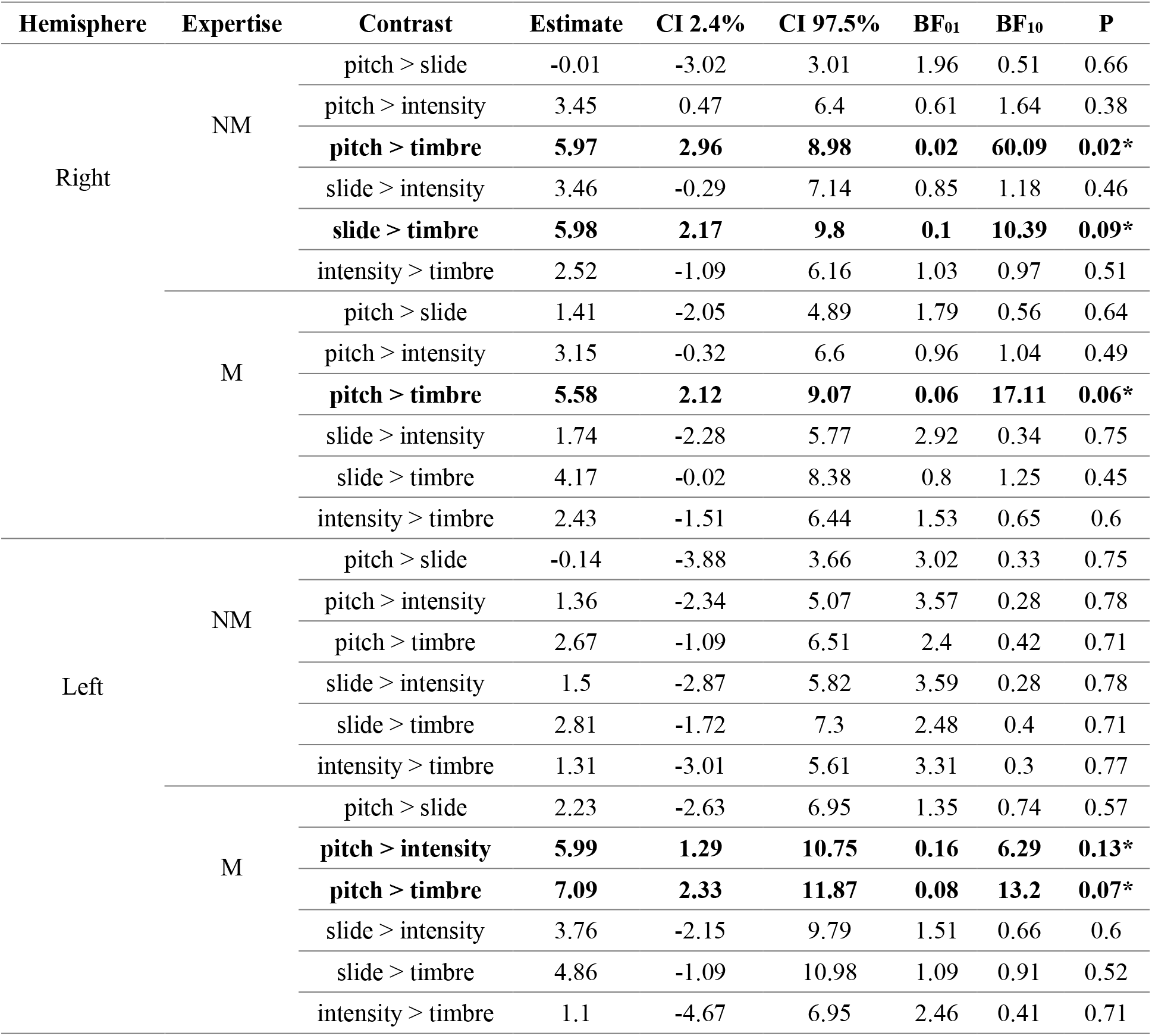
Pairwise Bayesian contrasts between features for entropy-related MMNm amplitude differences in each group and hemisphere. NM = non-musicians, M = musicians, CI = credible interval, BF_01_ = Bayes factor in favor of the null, BF_10_ = Bayes factor in favor of the alternative, P = posterior probability of the null. Contrasts with moderate or strong evidence for either the null hypothesis or the alternative hypothesis are highlighted in bold and marked with a star (*).

**Figure 5.**
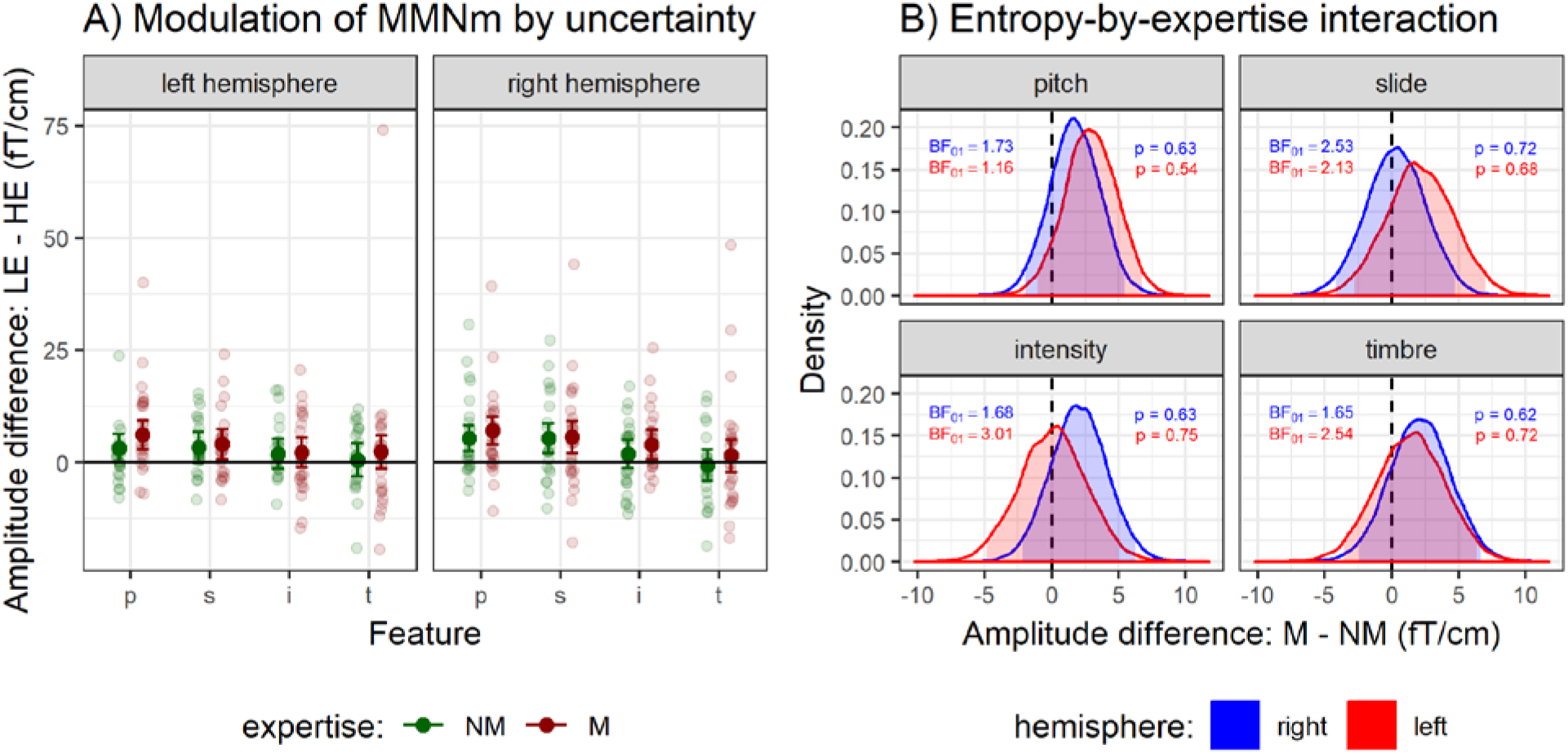
A) Bayesian estimates of the mean gradient MMNm amplitude differences between high-entropy (HE) and low entropy (LE) conditions for each group. Error bars represent 95% credible intervals. B) Posterior probability densities of the differences in the entropy effect between musicians and non-musicians (i.e. entropy-by-expertise interaction) for each hemisphere and feature. Shaded areas depict 95% credible intervals. NM = non-musicians, M = musicians, BF_M_ = Bayes factor in favor of the null, *p* = posterior probability of the null.

### 3.3. Source reconstruction

Neural generators of the MMNm were located in the surroundings of right and left auditory cortices, including both the posteromedial and anterolateral portions of Heschl’s gyrus (Figure 6). No prefrontal generators were observed, with the exception of the pitch MMNm for which there was a small source in the ventral part of the premotor cortex (BA6). Small clusters were also found for pitch in the somatosensory and parietal cortices, and for intensity in the parietal lobe around the perisylvian region. Regarding the entropy effect, the neural generators for pitch were located in the planum temporale anterior to the generator of the MMNm, whereas for slide a significant cluster was found in the right fusiform gyrus—an area involved in higher-order visual processing—, which could be related to spurious visual activity arising from watching the movie. For this reason, uncorrected values thresholded at .001 are shown for the entropy effect on the slide MMNm in the supplementary file 2, which includes clusters in the planum temporale.

**Figure 6.**
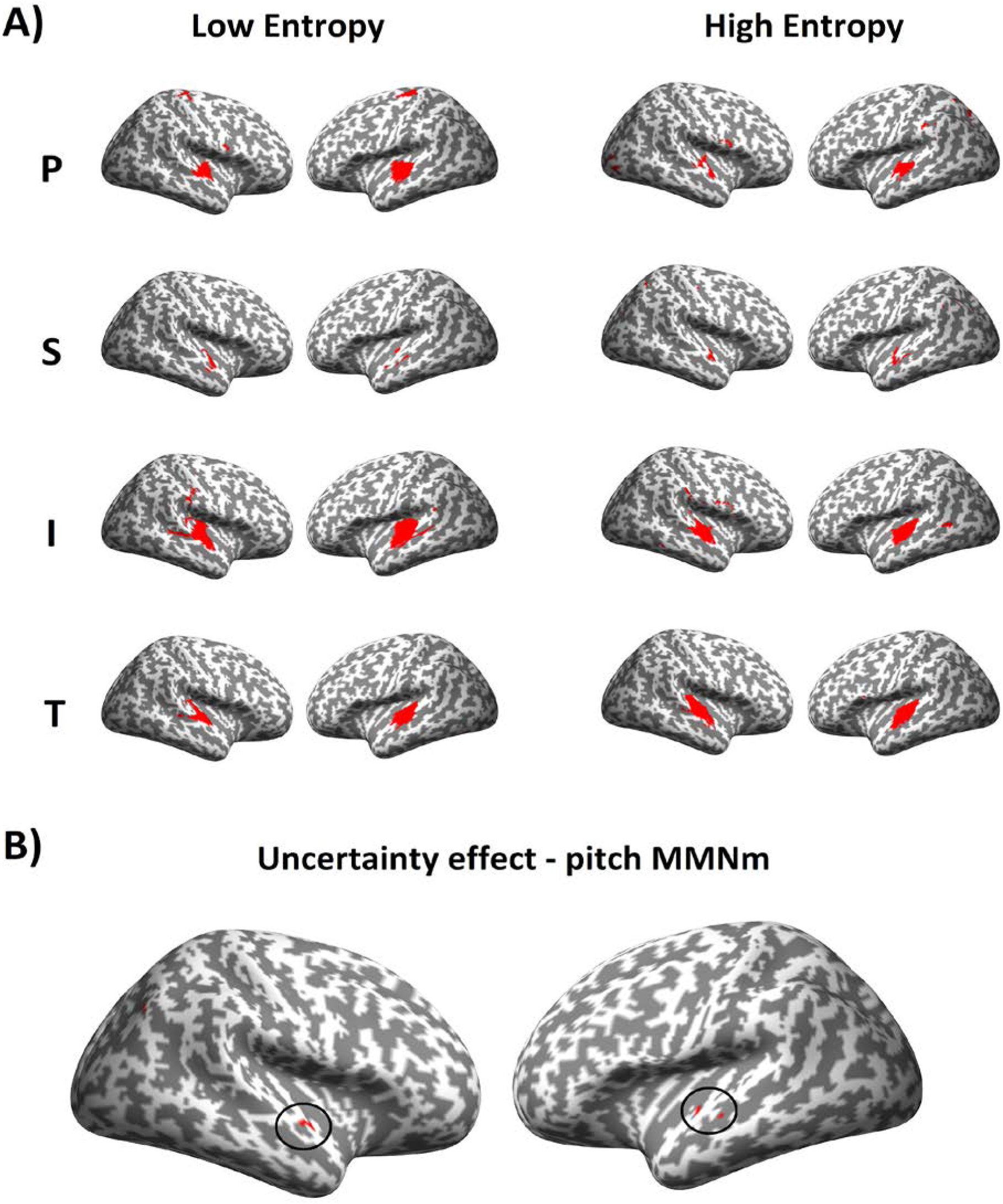
Statistical maps of the source reconstruction for A) the MMNm for each feature and condition and B) the effect of entropy on pitch MMNm responses. Clusters are thresholded at p <.05 after multiple-comparisons correction. Clusters for the entropy effect are marked with a circle. Participants from both groups (musicians and non-musicians) were included in the statistical tests. P = pitch, S = slide, I = intensity, T = timbre.

### 3.4. Behavioral experiment

Parameter estimates and data from the behavioral experiment are shown in Figure 7. Analyses of *d’* scores revealed that adding entropy as a categorical (*d1*) or continuous (d2) factor explained the data significantly better than an intercept-only (d0) model (Table 3). Furthermore, the comparisons *d1e-d1* and *d2e*-*d2* revealed a significant main effect of expertise, whereas the comparisons *d1i*-*d1e* and *d2i*-*d2e* were nonsignificant, thus failing to provide evidence for an entropy-by-expertise interaction. A contrast between the continuous (*d2e*) and the categorical model (*d1e*) was not significant (*χ^2^ = 1.53, p = .67*). The residuals of these two models were normally distributed. AIC values revealed a similar picture and slightly favored *d2e* over *d1e* as the winning model (Table 3). Bonferroni-corrected pairwise comparisons for the full model (*d1i*) showed significant differences between LE and the other four conditions for non-musicians. For musicians, however, the comparisons LE > IE1 and LE > IE2 were nonsignificant, whereas the contrasts LE > IE3, LE > HE, IE1 > IE2, IE1 > IE3 and IE1 > HE were significant. Finally, other comparisons such as IE1 > IE3, IE2 > HE and IE2 > HE, although nonsignificant, resulted in large effect sizes in both groups (table 4).

**Figure 7:**
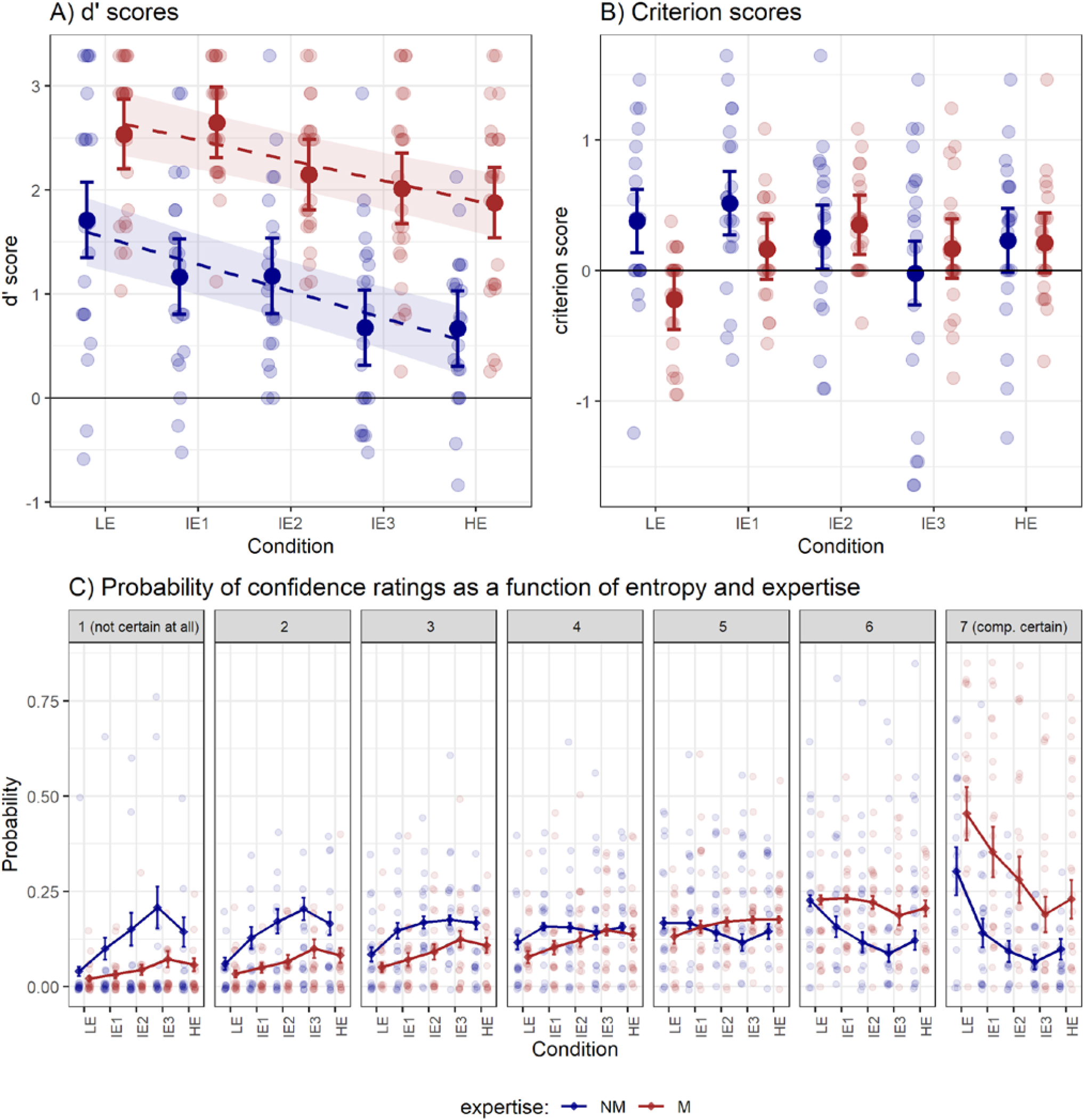
A) *d’* scores, B) *c* scores and C) confidence ratings—expressed as the probability of response for each confidence category. Note how the number of higher ratings (e.g. 7) tended to decrease, and the number of lower ratings (e.g. 1) tended to increase, with increasing entropy levels. Also note how musicians were more confident or certain overall. All parameter values were taken from maximum likelihood estimates. For *d’* scores, the slopes of the continuous (*d2i*) model are also plotted as dashed lines. Error bars and shaded areas represent 95% confidence intervals. M = musicians, NM = non-musicians, LE = low entropy, IE = intermediate entropy, HE = high entropy.

**Table 3.**
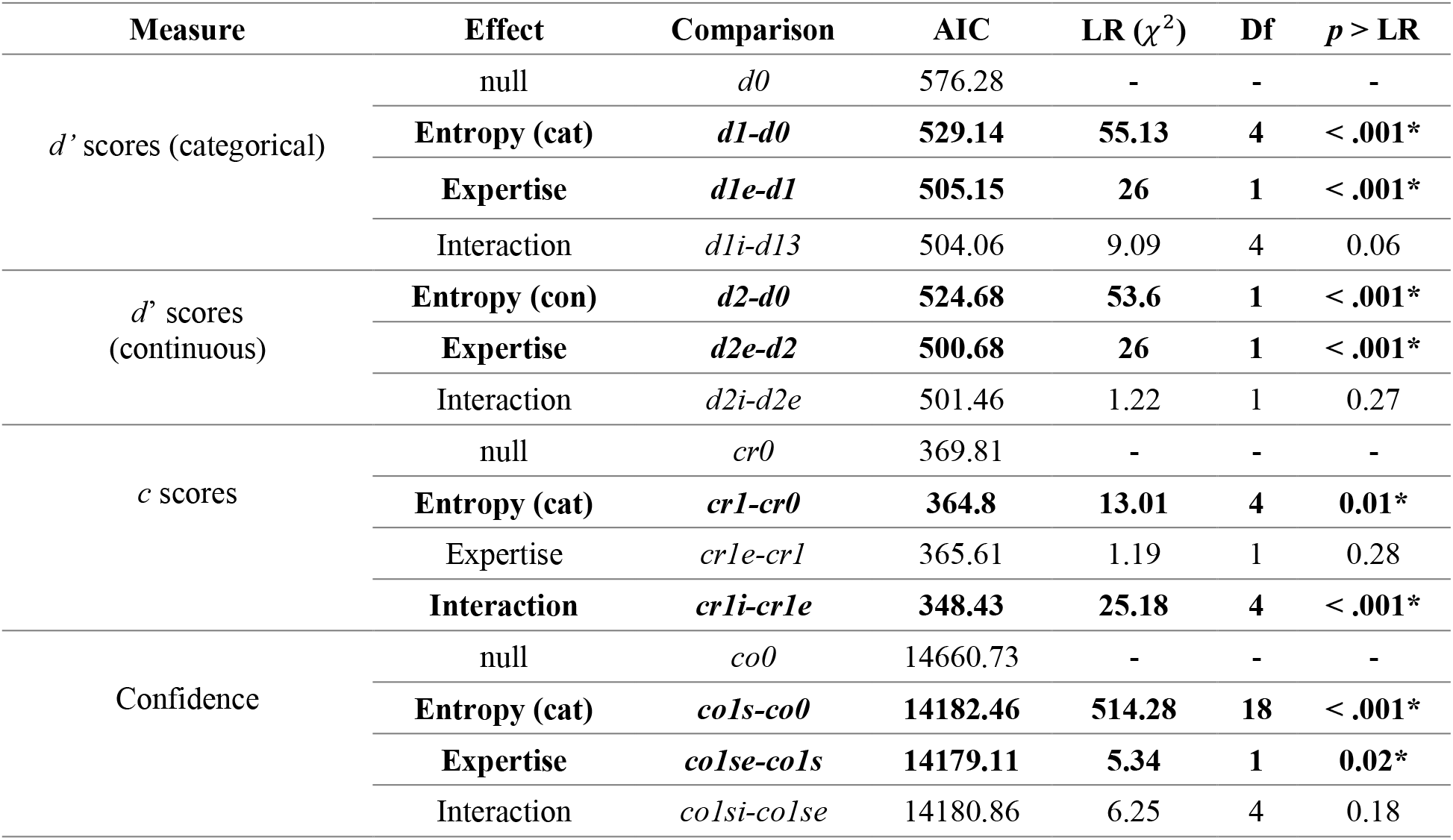
Likelihood ratio tests for all models in the behavioral experiment. AIC = Akaike Information Criterion, LR= Likelihood ratio, Df = difference in degrees of freedom, con = continuous, cat = categorical. Significant contrasts are highlighted in bold and marked with a star (*).

**Table 4.**
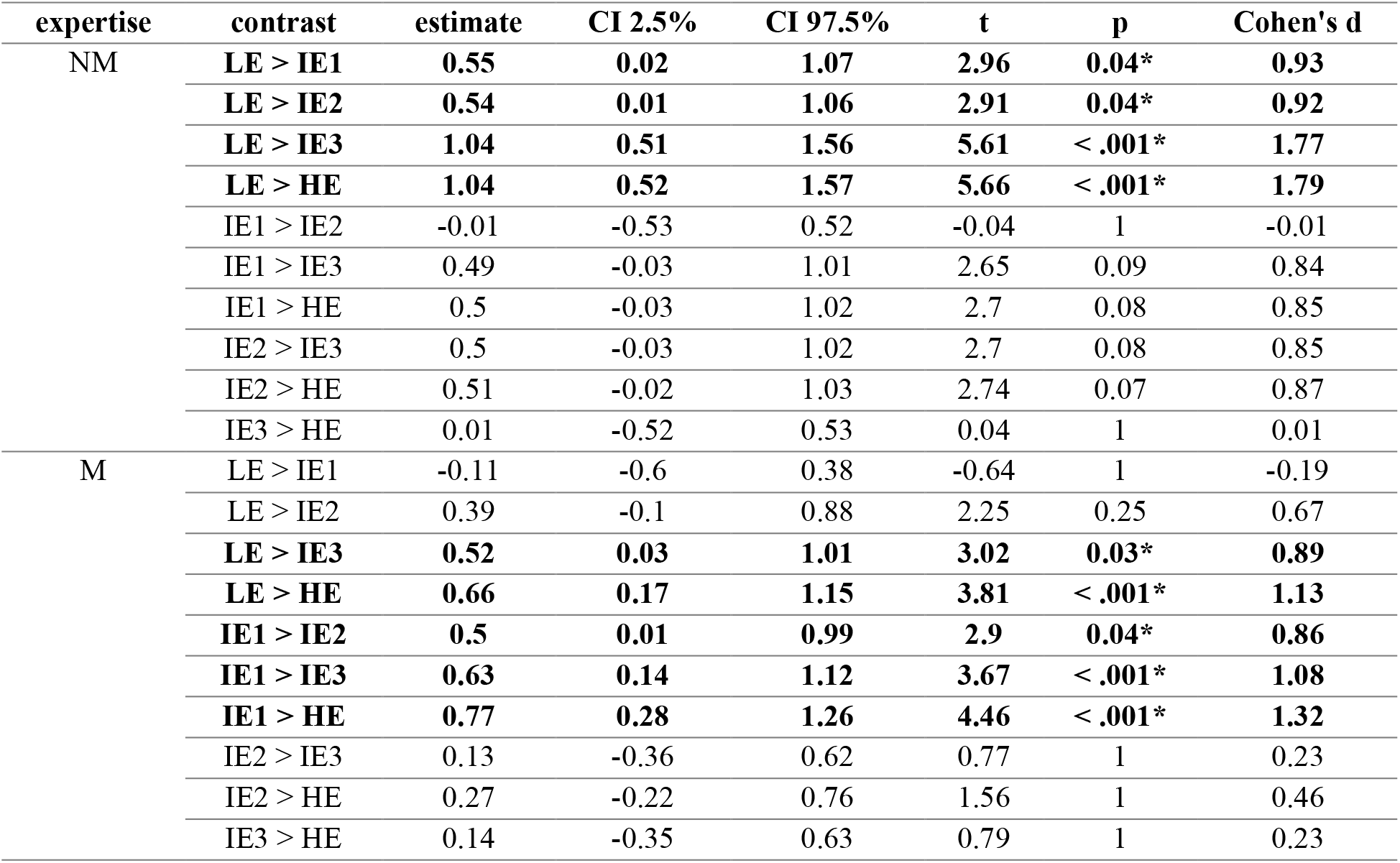
Bonferroni-corrected pairwise comparisons for d’-scores. Significant contrasts are highlighted in bold and marked with a star (*). CI = confidence interval, LE = low entropy, IE = intermediate entropy, HE = high entropy, M = musicians, NM = non-musicians.

| expertise | contrast | estimate | CI 2.5% | CI 97.5% | t | p | Cohen's d |
| --- | --- | --- | --- | --- | --- | --- | --- |
| NM | LE > IE1 | **0.55** | **0.02** | **1.07** | **2.96** | **0.04\*** | **0.93** |
|  | LE > IE2 | **0.54** | **0.01** | **1.06** | **2.91** | **0.04\*** | **0.92** |
|  | LE > IE3 | **1.04** | **0.51** | **1.56** | **5.61** | **< .001\*** | **1.77** |
|  | LE > HE | **1.04** | **0.52** | **1.57** | **5.66** | **< .001\*** | **1.79** |
|  | IE1 > IE2 | -0.01 | -0.53 | 0.52 | -0.04 | 1 | -0.01 |
|  | IE1 > IE3 | 0.49 | -0.03 | 1.01 | 2.65 | 0.09 | 0.84 |
|  | IE1 > HE | 0.5 | -0.03 | 1.02 | 2.7 | 0.08 | 0.85 |
|  | IE2 > IE3 | 0.5 | -0.03 | 1.02 | 2.7 | 0.08 | 0.85 |
|  | IE2 > HE | 0.51 | -0.02 | 1.03 | 2.74 | 0.07 | 0.87 |
|  | IE3 > HE | 0.01 | -0.52 | 0.53 | 0.04 | 1 | 0.01 |
| M | LE > IE1 | -0.11 | -0.6 | 0.38 | -0.64 | 1 | -0.19 |
|  | LE > IE2 | 0.39 | -0.1 | 0.88 | 2.25 | 0.25 | 0.67 |
|  | LE > IE3 | **0.52** | **0.03** | **1.01** | **3.02** | **0.03\*** | **0.89** |
|  | LE > HE | **0.66** | **0.17** | **1.15** | **3.81** | **< .001\*** | **1.13** |
|  | IE1 > IE2 | **0.5** | **0.01** | **0.99** | **2.9** | **0.04\*** | **0.86** |
|  | IE1 > IE3 | **0.63** | **0.14** | **1.12** | **3.67** | **< .001\*** | **1.08** |
|  | IE1 > HE | **0.77** | **0.28** | **1.26** | **4.46** | **< .001\*** | **1.32** |
|  | IE2 > IE3 | 0.13 | -0.36 | 0.62 | 0.77 | 1 | 0.23 |
|  | IE2 > HE | 0.27 | -0.22 | 0.76 | 1.56 | 1 | 0.46 |
|  | IE3 > HE | 0.14 | -0.35 | 0.63 | 0.79 | 1 | 0.23 |

Regarding Bayesian analyses, there was anecdotal/inconclusive evidence that the interaction terms were not different from zero, for both the *d1ib* and *d2ib* models (Figure 8). The only exception was the parameter for LE > IE1 in model *d1ib*, for which zero was located slightly to the left of the credible interval. An interaction in this case was about three times more likely than a null effect. This is in agreement with the likelihood ratio test between *d1i* and *d1e*, for which thep-value was close to the alpha threshold (table 3).

**Figure 8.**
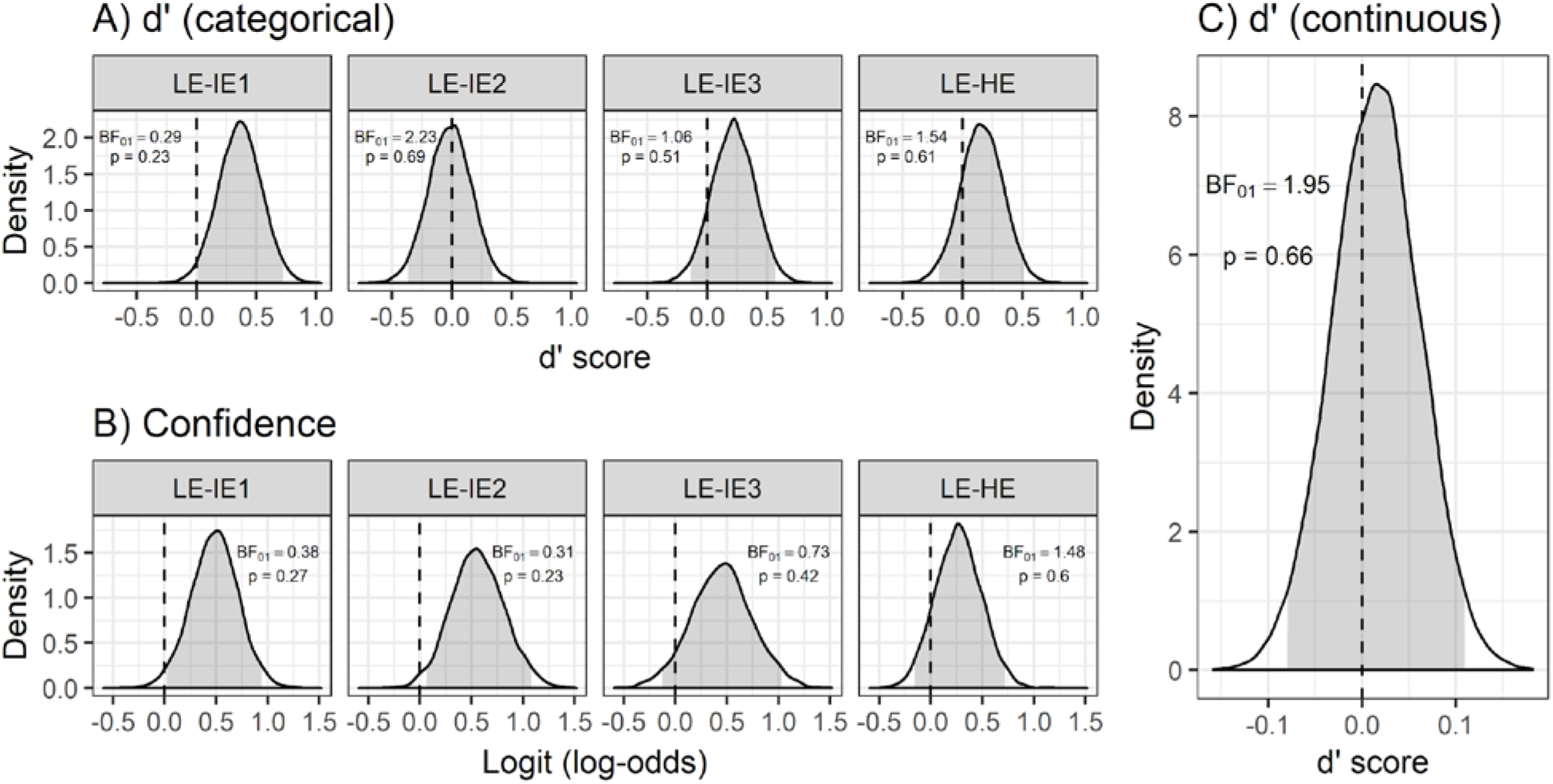
Bayesian posterior probability densities for the entropy-by-expertise interaction parameters—i.e. expertise-related modulation of the comparisons LE-IE1, LE-IE2, LE-IE3, LE-HE and the slope of the continuous model. Densities for (A) categorical and (C) continuous mixed models of d’ scores, as well as (B) a cumulative-link mixed model of confidence ratings are displayed. Shaded areas represent 95% credible intervals. Since accuracy and confidence generally decreased with higher entropy levels, positive parameter values indicate smaller differences between conditions for musicians compared to non-musicians. BF_01_ = Bayes factor in favor of the null, p = posterior probability of the null, LE = low entropy, IE = intermediate entropy, HE = high entropy.

An analysis of *c* scores revealed a main effect of entropy and an interaction between entropy and expertise (Table 3). The mean *c* score for both groups was positive, thus indicating a mild bias towards missing the targets. The bias changed between conditions following different patterns for each group, as revealed in the pairwise contrasts. Concretely, for non-musicians there was a significant difference in the contrasts LE > IE3 and IE1 > IE3, whereas for musicians *c* scores were significantly higher for LE than the other four conditions (supplementary materials 3). When contrasting musicians and non-musicians for each condition separately, the nature of the interaction was clearer. For LE and IE1, the bias was significantly lower for musicians than non-musicians and even became negative in the case of LE (Table 6, Figure 7).

**Table 6.** Bonferroni-corrected contrasts of c scores between musicians and non-musicians for each condition. Significant contrasts are highlighted in bold and marked with a star (*). CI = confidence interval, LE = low entropy, IE = intermediate entropy, HE = high entropy, M = musicians, NM = non-musicians.

| condition | contrast | estimate | CI 2.5% | CI 97.5% | t | p | Cohen's d |
| --- | --- | --- | --- | --- | --- | --- | --- |
| LE | NM - M | **0.6** | **0.26** | **0.94** | **3.46** | **< .001\*** | **1.41** |
| IE1 | NM - M | **0.35** | **0.01** | **0.7** | **2.05** | **0.04\*** | **0.84** |
| IE2 | NM - M | -0.1 | -0.44 | 0.25 | -0.55 | 0.58 | -0.22 |
| IE3 | NM - M | -0.19 | -0.53 | 0.16 | -1.08 | 0.28 | -0.44 |
| HE | NM - M | 0.02 | -0.32 | 0.36 | 0.1 | 0.92 | 0.04 |

Regarding confidence ratings, there were main effects of entropy and expertise, as revealed by the contrasts *co1s-co0* and *colse-cols*, respectively (Table 3). Adding an interaction term (*co1si*) did not explain the data significantly better. AIC values suggested *co1se* as the winning model. Pairwise comparisons revealed significant differences between LE and the other four conditions and the contrast IE1 > IE3 for non-musicians (Table 5). For musicians, there was not a significant difference for the contrast LE > IE1 but for contrasts IE1 > LE2, IE1 > LE3, IE > HE, IE1 > IE3, IE1 > HE and IE2 > IE3. Bayesian analyses suggested moderate evidence for an interaction in the case of the LE-IE1 and LE-IE2 slopes and inconclusive evidence in favor of the null for the LE-IE3 and LE-HE parameters (Figure 8).

**Table 5.** Bonferroni-corrected pairwise comparisons for confidence ratings. Significant contrasts are highlighted in bold and marked with a star (*). CI = confidence interval, LE = low entropy, IE = intermediate entropy, HE = high entropy, M = musicians, NM = non-musicians.

| expertise | contrast | odds ratio | CI 2.5% | CI 97.5% | z | p |
| --- | --- | --- | --- | --- | --- | --- |
| NM | LE > IE1 | **2.64** | **1.59** | **4.38** | **5.39** | **< .001\*** |
|  | LE > IE2 | **4.24** | **2.09** | **8.59** | **5.74** | **< .001\*** |
|  | LE > IE3 | **6.23** | **2.69** | **14.44** | **6.11** | **< .001\*** |
|  | LE > HE | **3.99** | **2.19** | **7.25** | **6.49** | **< .001\*** |
|  | IE1 > IE2 | 1.6 | 0.84 | 3.08 | 2.03 | 0.42 |
|  | IE1 > IE3 | **2.36** | **1.04** | **5.35** | **2.94** | **0.03\*** |
|  | IE1 > HE | 1.51 | 0.82 | 2.79 | 1.88 | 0.6 |
|  | IE2 > IE3 | 1.47 | 0.96 | 2.24 | 2.57 | 0.1 |
|  | IE2 > HE | 0.94 | 0.65 | 1.36 | -0.46 | 1 |
|  | IE3 > HE | 0.64 | 0.4 | 1.01 | -2.74 | 0.06 |
| M | LE > IE1 | 1.53 | 0.92 | 2.52 | 2.36 | 0.18 |
|  | LE > IE2 | **2.14** | **1.08** | **4.21** | **3.14** | **0.02\*** |
|  | LE > IE3 | **3.56** | **1.6** | **7.92** | **4.46** | **< .001\*** |
|  | LE > HE | **2.79** | **1.56** | **4.99** | **4.96** | **< .001\*** |
|  | IE1 > IE2 | 1.4 | 0.75 | 2.62 | 1.5 | 1 |
|  | IE1 > IE3 | **2.33** | **1.07** | **5.08** | **3.06** | **0.02\*** |
|  | IE1 > HE | **1.83** | **1.01** | **3.31** | **2.86** | **0.04\*** |
|  | IE2 > IE3 | **1.67** | **1.1** | **2.52** | **3.48** | **0.01\*** |
|  | IE2 > HE | 1.31 | 0.9 | 1.89 | 2.02 | 0.43 |
|  | IE3 > HE | 0.78 | 0.5 | 1.22 | -1.54 | 1 |

## 4. Discussion

In the present work, we show that the reduction of prediction error responses by predictive uncertainty (Quiroga-Martinez et al., 2019) is also found in musically trained participants. This indicates that the effect is robust and is present across listeners with different degrees of musical expertise. Moreover, while musicians had larger MMNm responses to pitch and slide deviants, there was no evidence for an entropy-by-expertise interaction that would indicate a less pronounced effect of uncertainty for musical experts.

### 4.1. Expertise-related effects

Across different degrees of melodic entropy, musicians were more accurate and confident in detecting pitch deviations, and had larger pitch and slide MMNm responses in the MEG experiment. This extends previous findings showing larger MMNm responses—especially for pitch-related deviants—in musicians than non-musicians (Brattico et al., 2009; Fujioka et al., 2004; Koelsch et al., 1999; Putkinen et al., 2014; Tervaniemi et al., 2014; Vuust et al., 2012, 2005). Since this indicates an enhancement of auditory discrimination skills and could be framed as an increase in predictive precision, it is rather surprising that the effect of contextual uncertainty on prediction error is not significantly different between the two groups.

What these results suggest is that expertise-driven and stimulus-driven changes in predictive precision are dissociable and independent. Thus, intensive musical training might sharpen the long-term representation of in-tune musical pitch categories and therefore facilitate the detection of pitch deviants. This would result in higher baseline levels of deviance detection and confidence as well as larger baseline MMNm amplitudes for musicians. In contrast, the uncertainty of the current stimulus would be inferred dynamically from its local statistics, leading to a short-term modulation of prediction error that is independent from long-term expertise, hence explaining the additive effects observed in both the MEG and behavioral experiments.

The observed pattern of results is surprising also with regard to behavioral studies in which musicians gave significantly higher unexpectedness ratings to melodic continuations than non-musicians, in contexts with low but not high entropy (Hansen & Pearce, 2014; Hansen et al., 2016). This finding has been taken to reflect a better ability of musicians to distinguish between low-and high-entropy contexts. Note that these results would predict a larger effect of entropy in musical experts, which is the opposite of what we hypothesized, but for which there was no evidence in our data either. It has to be noted, though, that the type of unexpected tones that we used was different from the one reported in those experiments. Here, surprising tones corresponded to out-of-tune deviants, whereas in the behavioral studies unexpectedness judgements were made on plausible in-tune melodic continuations. Furthermore, the effect of entropy was reported for expected and unexpected tones combined, whereas here we only employed tones that were highly unexpected. These discrepancies point to future research addressing the effect of entropy on the neural responses to in-tune compared to out-of-tune surprising tones, and to expected and unexpected tones separately.

Bayesian estimation allowed us to evaluate the relative evidence for the null and alternative hypotheses. For the change in MMNm amplitude between conditions, the parameter estimates of the difference between groups generally had small mean values (Figure 5b), indicating a rather small or absent modulation of expertise. However, while all credible intervals contained zero, they were also uncertain, spanning a rather broad amplitude range. This is reflected in Bayes factors, which were inconclusive, and therefore not much can be said about the null hypothesis.

In the behavioral experiment, the picture was slightly different. For d’-scores, there was moderate evidence that the difference between LE and IE1 was reduced in musical experts. Evidence for other interaction terms, including LE-IE3, LE-HE and the slope for the continuous model, although inconclusive, suggested that the effect of entropy was slightly less pronounced for musicians. A similar pattern was observed for the confidence ratings. Therefore, although likelihood ratio tests were non-significant for the interactions, Bayesian analyses provided some evidence for a group difference, at least for some interaction parameters.

Based on this, it would be tempting to conclude that there is evidence for our hypothesis at the behavioral level. However, these results might as well arise from a ceiling effect in musicians’ *d’* scores and confidence ratings that would reduce differences between LE and IE1 or LE and IE2, compared to the non-musicians’ noticeable differences between these conditions. The distribution of individual data points in Figure 7 suggests that this might be the case. Taking into consideration the results of the MEG experiment, the generally inconclusive Bayes factors and the possibility of a ceiling effect, it is fair to remain skeptical about the presence of an entropy-by-expertise interaction when the two experiments are considered together.

### 4.2. Feature-specific effects

One intriguing finding in Quiroga-Martinez et al. (2019) was that the effect of predictive uncertainty on prediction error was restricted to pitch-related deviants (out-of-tune tones and pitch glides). This was interpreted as suggesting a feature-selective effect, given that uncertainty was manipulated in the pitch dimension only, while uncertainty in other dimensions such as timbre and intensity was kept constant. In the current work, this result was replicated in musicians, although with a slightly different pattern of differences. In non-musicians, larger entropy effects were observed for pitch and slide deviants, compared to intensity and timbre deviants, in the right but not the left hemisphere. For musicians, larger entropy effects were found for pitch deviants when compared with timbre deviants in both hemispheres, and when compared with intensity deviants in the left hemisphere.

Care should be taken not to overinterpret these potential expertise-related differences, until they have been shown in direct group comparisons. However, attention should be paid to the intensity MMNm because, for musicians, a small yet significant difference between LE and HE contexts was found in the cluster-based permutation analyses, which in turn resulted in a significant main effect of entropy for this feature. Note that, for non-musicians, there was already a hint of such a difference. Therefore, it seems that intensity prediction errors are also somewhat affected by the pitch entropy of the melodies, something that challenges the proposed feature-selectivity.

These results suggest that uncertainty is mainly feature-selective, but has a residual effect on predictive processing in other features as well. However, there might be two confounding factors here. First, the perception of loudness changes with pitch height (Suzuki, Møller, Ozawa, & Takeshima, 2003). In that case, the slightly different pitch distributions in HE and LE conditions (see Supplementary figure 1 in Quiroga-Martinez, et al., 2019) might have made the loudness violation slightly more or less salient for different conditions. The second confound could be the baseline salience of the deviants, which might have differed between features. For example, a very strong timbre violation might have been less affected by entropy than a less strong intensity violation or an even weaker pitch violation, thus yielding the observed feature-specific patterns.

Another interesting feature-wise finding is that musicians had larger MMNm amplitudes for pitch and slide but not timbre or intensity compared to non-musicians. This is consistent with the literature, in which larger amplitudes have been consistently found for pitch-related deviants in musicians but less so for other features (Putkinen et al., 2014; Tervaniemi et al., 2014). This might reflect a focus on pitch discrimination as a core ability for musical experts and the fact that musical pitch is organized in rich multidimensional cognitive systems (Krumhansl, 1990), which is not the case for intensity or timbre.

### 4.3. Source reconstruction

As expected from the literature (Deouell, 2007), neural generators of the MMNm were located in primary and secondary bilateral auditory cortices. No prefrontal generators were observed, with the exception of the pitch MMNm for which there seemed to be a small source in the ventral part of BA6. This, however, could be caused by leakage of the temporal source. The location of the entropy effect for the pitch MMNm, which was anterior to the primary source, suggests that entropy affected the passing of prediction error responses from primary to secondary auditory cortex. This is consistent with predictive processing theories (Clark, 2016; Feldman & Friston, 2010; Hohwy, 2013) that suggest an uncertainty-driven reduction in the gain of prediction error responses, which prevents them from driving inference and learning at higher levels of the cortical hierarchy (Griffiths & Warren, 2002). Moreover, this is partly consistent with results reported by Southwell and Chait (2018), who found reduced anterior temporal responses to deviant sounds in contexts with high as compared to low uncertainty. Note that other prefrontal sources were also found in that study which were not detected here. However, their results were based on EEG source reconstructions with no individual anatomical images, which may explain the differences in the sources found in the two studies.

### 4.4. Behavioral experiment

As mentioned above, the effect of entropy on accuracy and confidence scores was present in musicians as well. There was also a main effect of expertise in which musicians were better and more confident at discriminating the deviants, but there was no conclusive evidence for an entropy-by-expertise interaction. Thus, behavioral measures were in agreement with the outcomes of the MEG experiment. Apart from these findings, two results deserve attention. First, pairwise differences were found between HE and IE1, and between intermediate conditions (e.g. IE1 and IE3) in both groups. This corroborates the finding that any of the two sources of uncertainty manipulated in the experiment, namely pitch-alphabet size and stimulus repetitiveness, can modulate prediction error separately. Second, *c* scores revealed a reduced bias in musicians for LE and IE1 conditions, indicating a higher rate of hits only for categories with the highest precision. Interestingly, for LE stimuli, some musicians systematically reported deviants when there were none, which suggests that the expectancy of a deviation might have occasionally induced the illusion of a mistuning.

### 4.5. Limitations and future directions

Since we used the same methods as in Quiroga-Martinez et al. (2019), the limitations already discussed in that work also apply to the current report. Briefly, these include the impossibility of disentangling the contribution of pitch-alphabet size and repetitiveness to the modulation of the MMNm; the different repetition rates of individual melodies in different conditions, which might have created different veridical expectations during stimulation; the difference in the distribution of pitches between conditions in the MEG experiment and its possible implications for the pitch MMNm; the measurement of uncertainty at the context level rather than on a note-by-note basis; the unusual listening situation—i.e. participants listening passively while watching a silent movie—which limits the generality of the findings; the rather artificial auditory context—even though our stimuli are much more realistic than in most MMN research; and the lack of a preregistration of our hypothesis and analysis plan—something that is partly overcome by the fact that we have now replicated our main findings and have openly shared materials, code and data.

Despite these limitations, our work provides further evidence for the effect of uncertainty—or precision— on prediction error, which is consistent with an increasing number of empirical findings (Garrido et al., 2013; Hsu et al., 2015; Lumaca et al., 2019; Sedley et al., 2016; Sohoglu & Chait, 2016; Southwell & Chait, 2018), theories of predictive processing and models of music perception (Clark, 2016; Feldman & Friston, 2010; Hohwy, 2013; Ross & Hansen, 2016; Vuust et al., 2018). Furthermore, our findings confirm that MMNm responses can be reliably recorded in realistic paradigms where sounds constantly change, which constitutes a methodological improvement on existing approaches.

Consequently, we hereby open the possibility of addressing questions about predictive processing and predictive uncertainty in more realistic and complex auditory environments. One possible future direction in this regard would be to elucidate where and how the modulation of prediction error takes place in the auditory frontotemporal network. Specifically, it would be interesting to address whether the precision-weighting effect arises from top-down or intrinsic connectivity, and whether neuromodulation plays a role (Auksztulewicz et al., 2018). Paradigms similar to the one presented here could be used in combination with connectivity measures, such as dynamic causal modelling (Moran, Pinotsis, & Friston, 2013), and intracranial recordings (e.g. Omigie et al., 2019) to address these questions. Moreover, for music research, methods such as these could be very informative about the nature of musical knowledge and musical expectations and how these are represented in the cortical hierarchy. Relatedly, this line of research could inform musical aesthetics, given that some musical styles exploit uncertainty as an artistic resource (Mencke, Omigie, Wald-Fuhrmann, & Brattico, 2019). Finally, the use of more realistic stimuli could help us understand how different types of musical stimuli (e.g. different styles) are processed by listeners of different backgrounds, something that we have started to address here with musical experts, but that could be extended, for example, to listeners from different cultures or with instrument-specific expertise.

## Conclusion

In the present study we have shown that pitch prediction error responses in musical experts—as indexed by MMNm responses, accuracy scores, and confidence ratings—are reduced by pitch predictive uncertainty when listening to relatively complex and realistic musical stimuli. This suggests that our previous findings in non-musicians are robust and replicable and provides further support for theories of predictive processing which propose that neural responses to surprising stimuli are modulated by predictive uncertainty. Furthermore, our results show that, while musicians have generally larger prediction error responses, the uncertainty effect does not substantially change with expertise, thus pointing to separate long-term and short-term mechanisms of precision modulation. Overall, our work demonstrates that music, as a rich and multifaceted auditory signal, is an ideal means to improve our understanding of uncertainty and predictive processing in the brain.

## Acknowledgments

We wish to thank the project initiation group, namely Christopher Bailey, Torben Lund and Dora Grauballe, for their help with setting up the experiments. We also thank Nader Sedghi, Massimo Lumaca, Giulia Donati, Ulrika Varankaite, Giulio Carraturo, Riccardo Proietti, and Claudia Iorio for assistance during MEG recordings. We are indebted as well to the group of Italian trainees fromIISS Simone-Morea, Conversano, who helped with the behavioral experiment. Finally, we thank Hella Kastbjerg for checking the English language of this manuscript. The Center for Music in the Brain is funded by the Danish National Research Foundation (DNRF 117), which did not have any influence on the scientific content of this article.

## CRediT authorship contribution statement

David R. Quiroga-Martinez: Conceptualization, Methodology, Software, Formal analysis, Data curation, Writing - original draft, Visualization, Investigation. Niels C. Hansen: Conceptualization, Methodology, Writing - original draft. Andreas Højlund: Conceptualization, Methodology, Software, Writing - original draft, Supervision. Marcus T. Pearce: Software, Formal analysis, Writing - original draft. Elvira Brattico: Conceptualization, Supervision, Writing - original draft. Peter Vuust: Conceptualization, Methodology, Supervision, Writing - original draft.

