## Supplementary file 1 for "Musical prediction error responses similarly reduced by predictive uncertainty in musicians and non-musicians"

Standard and deviant Event Related Fields for musicians and non-musicians in high- and low-entropy contexts. The displayed activity corresponds to the average of the four right temporal combined gradiometers with the largest amplitude (channels 1342-1343, 1312-1313, 1322-1323 and 1332-1333). Gray lines depict individual MMNm responses. Shaded gray areas indicate 95% confidence intervals. Dashed vertical lines mark tone onsets. For descriptive purposes, green horizontal lines indicate when this difference was significant, according to the permutation tests. Note, however, that this is not an accurate estimate of the true extent of the effect

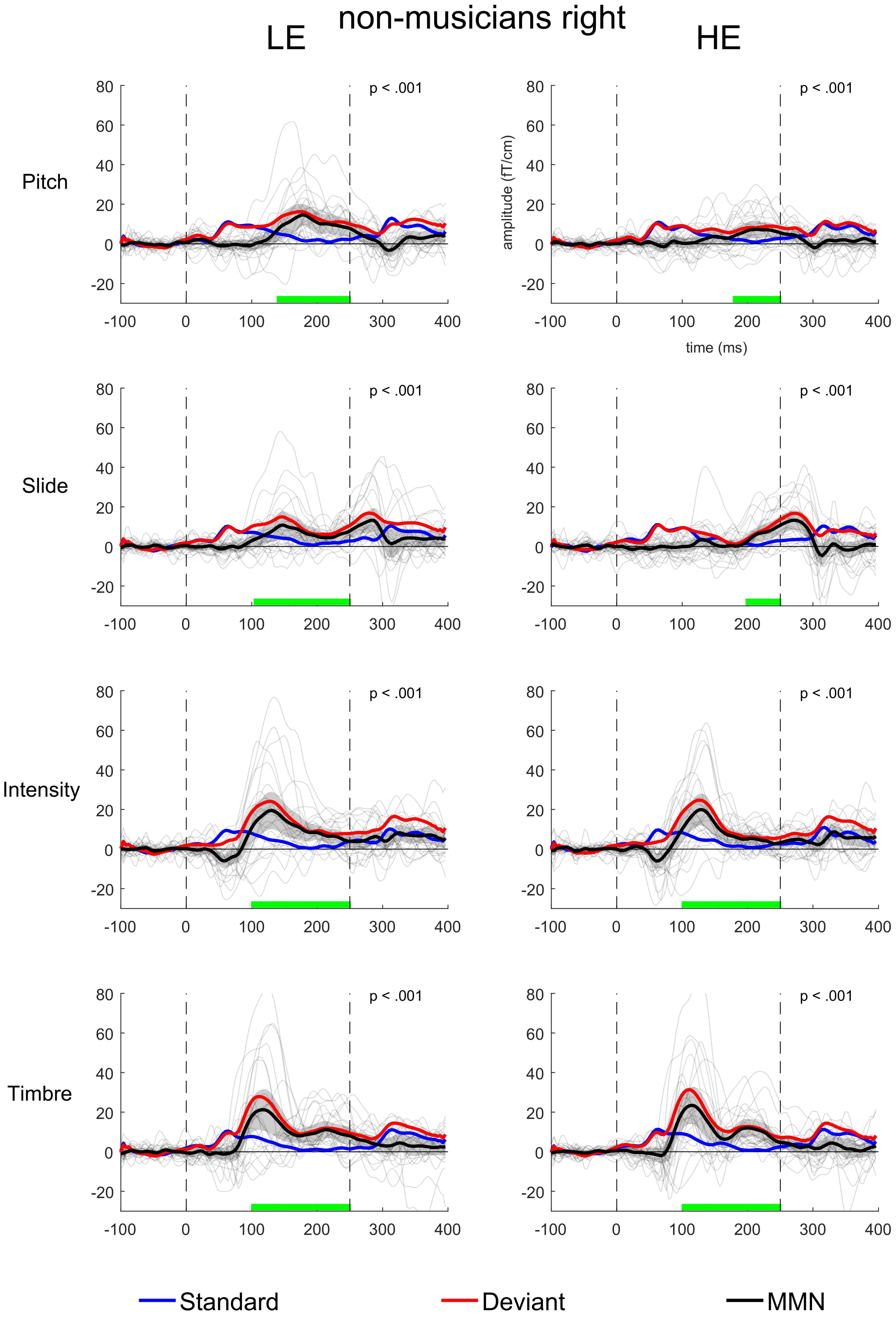
non-musicians right
LE
HE
Pitch
Slide
Intensity
Timbre
p < .001
amplitude (fT/cm)
time (ms)
Standard
Deviant
MMN

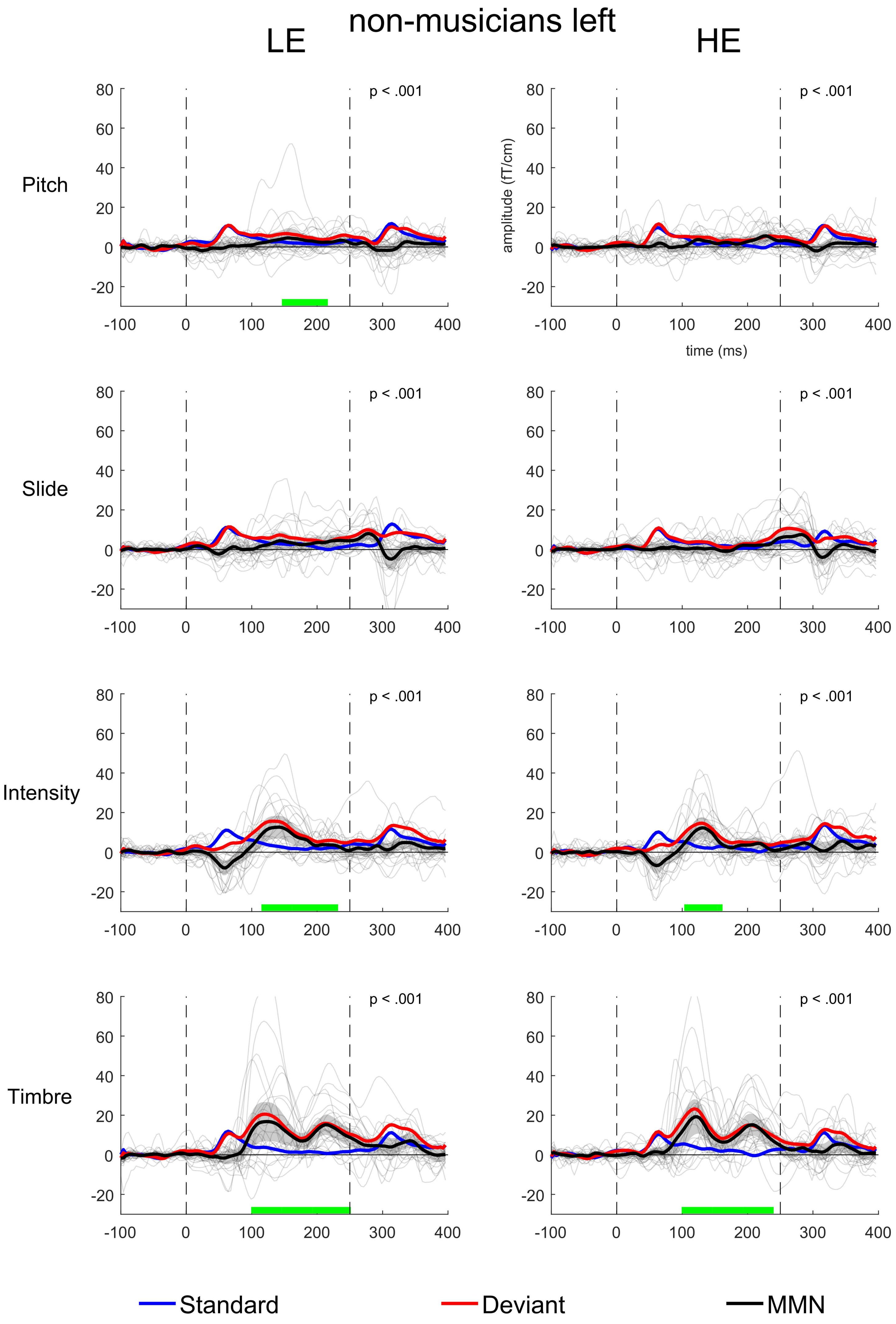
non-musicians left
LE
HE
Pitch
Slide
Intensity
Timbre
p < .001
amplitude (fT/cm)
time (ms)
Standard
Deviant
MMN

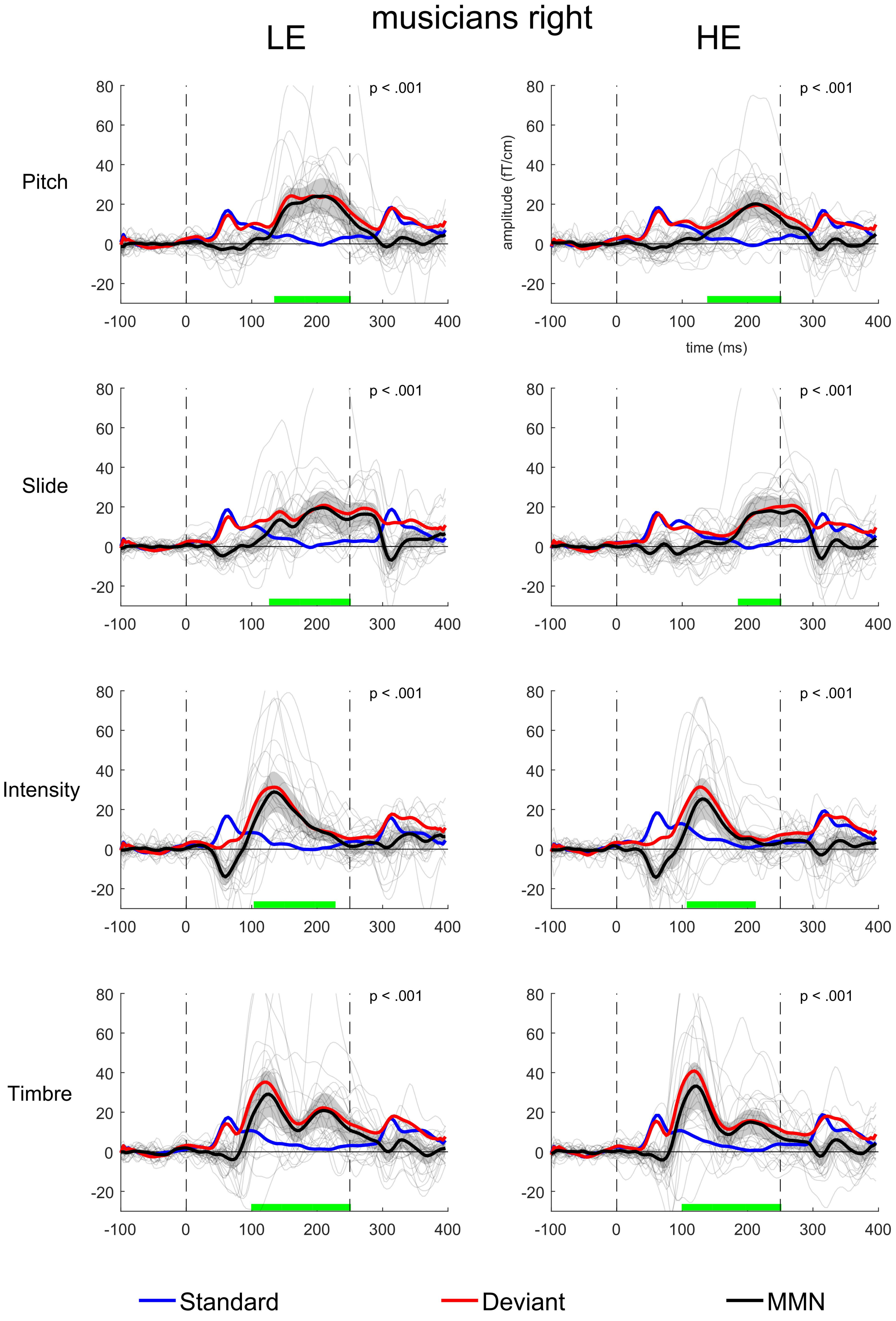
musicians right
LE
HE
Pitch
Slide
Intensity
Timbre
p < .001
amplitude (fT/cm)
time (ms)
Standard
Deviant
MMN

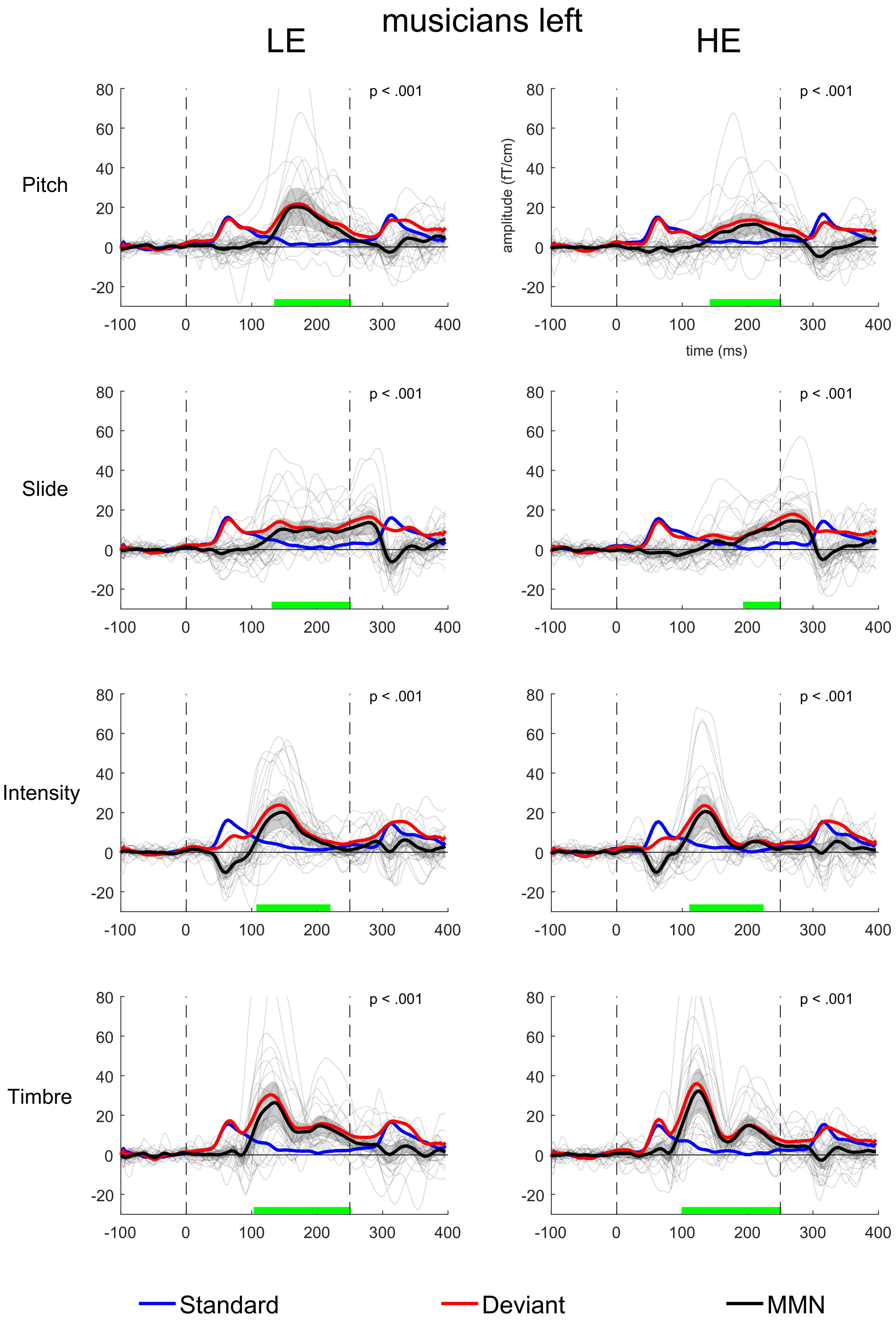

musicians left
LE
HE
Pitch
Slide
Intensity
Timbre
p < .001
amplitude (fT/cm)
time (ms)
Standard
Deviant
MMN
