## Supplementary figures and images for "Musical prediction error responses similarly reduced by predictive uncertainty in musicians and non-musicians"

### Suplementary file 2

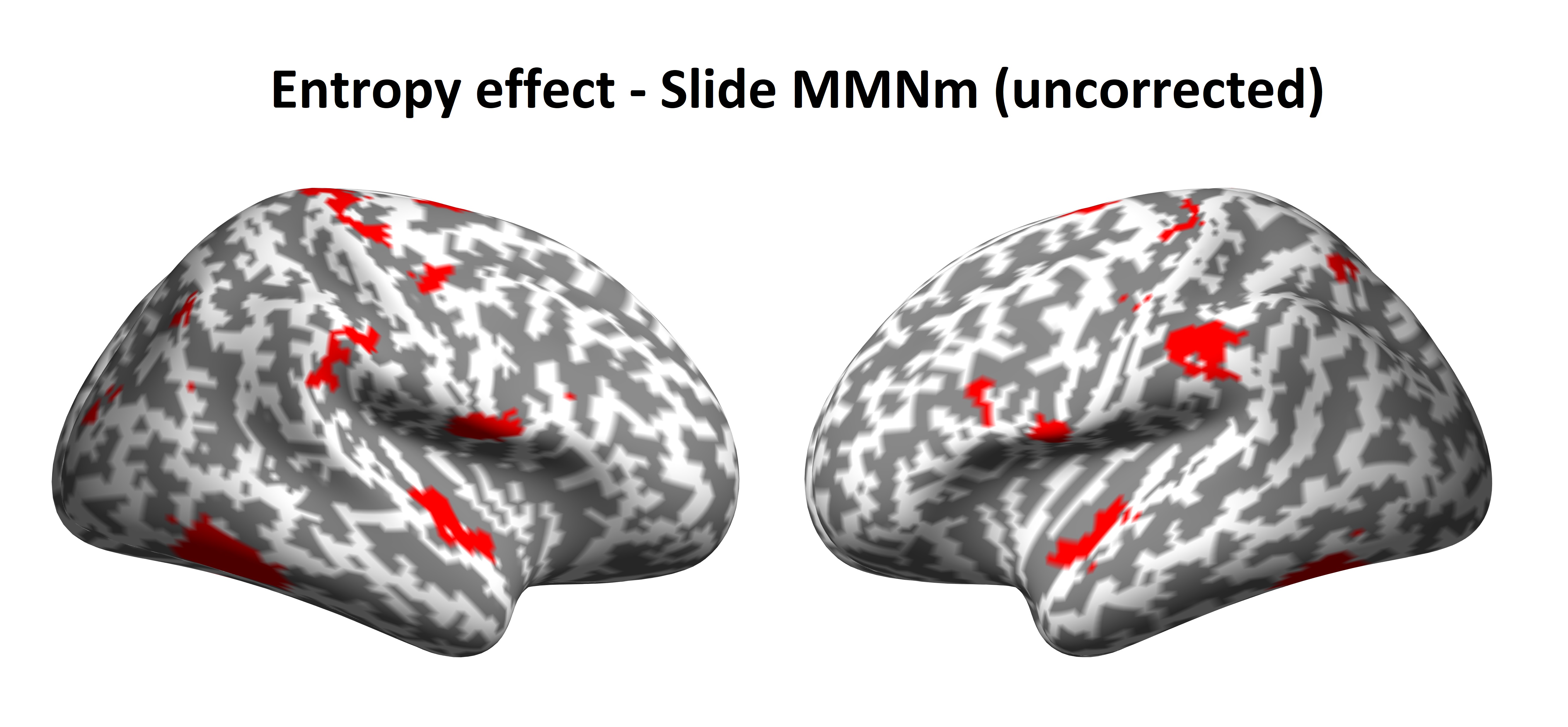
